# Complete genome sequencing of *Acinetobacter baumannii* AC1633 and *Acinetobacter nosocomialis* AC1530 unveils a large multidrug resistant plasmid encoding the NDM-1 and OXA-58 carbapenemases

**DOI:** 10.1101/2020.10.08.331108

**Authors:** Ahmed Ghazi Alattraqchi, Farahiyah Mohd. Rani, Nor Iza A. Rahman, Salwani Ismail, David W. Cleary, Stuart C. Clarke, Chew Chieng Yeo

**Affiliations:** Faculty of Medicine, Universiti Sultan Zainal Abidin, Kuala Terengganu, Terengganu, Malaysia; Faculty of Medicine and Institute for Life Sciences, University of Southampton, Southampton, UK; NIHR Southampton Biomedical Research Centre, University Hospital Southampton NHS Trust, Southampton, UK; Global Health Research Institute, University of Southampton, Southampton, UK; School of Postgraduate Studies, International Medical University, Kuala Lumpur, Malaysia; Centre for Translational Research, IMU Institute for Research, Development and Innovation (IRDI), Kuala Lumpur, Malaysia

**Author notes:** Corresponding author: Chew Chieng Yeo, PhD., Faculty of Medicine, Universiti Sultan Zainal Abidin (Medical Campus), Jalan Sultan Mahmud, 20400 Kuala Terengganu, Terengganu, MALAYSIA. Ahmed Ghazi Alattraqchi and Farhaiyah Mohd. Rani contributed equally to this work. Author order was determined by alphabetical order.

**Keywords:** NDM-1, OXA-58, carbapenem resistance, Tn125, *dif* modules, *Acinetobacter baumannii*, *Acinetobacter nosocomialis*, plasmids

## Abstract

Carbapenem-resistant *Acinetobacter* spp. are considered priority drug-resistant human pathogenic bacteria. The genomes of two carbapenem-resistant *Acinetobacter* spp. clinical isolates obtained from the same tertiary hospital in Terengganu, Malaysia, namely *A. baumannii* AC1633 and *A. nosocomialis* AC1530, were sequenced. Both isolates were found to harbor the carbapenemase genes *bla*_NDM-1_ and *bla*_OXA-58_ in a large (ca. 170 kb) plasmid designated pAC1633-1 and pAC1530, respectively, that also encodes genes that confer resistance to aminoglycosides, sulfonamides, and macrolides. The two plasmids were almost identical except for the insertion of IS*Aba11* and an IS*4* family element in pAC1633-1, and IS*Aba11* along with *relBE* toxin-antitoxin genes flanked by inversely orientated p*dif* (XerC/XerD) recombination sites in pAC1530. The *bla*_NDM-1_ gene was encoded in a *Tn125* composite transposon structure flanked by IS*Aba125* whereas *bla*_OXA-58_ was flanked by IS*Aba11* and IS*Aba3* downstream and a partial IS*Aba3* element upstream within a p*dif* module. The presence of conjugative genes in plasmids pAC1633-1/pAC1530 and their discovery in two distinct species of *Acinetobacter* from the same hospital are suggestive of conjugative transfer but mating experiments failed to demonstrate transmissibility under standard laboratory conditions. Comparative sequence analysis strongly inferred that pAC1633-1/pAC1530 was derived from two separate plasmids in an IS*1006*-mediated recombination or transposition event. *A. baumannii* AC1633 also harbored three other plasmids designated pAC1633-2, pAC1633-3 and pAC1633-4. Both pAC1633-3 and pAC1633-4 are cryptic plasmids whereas pAC1633-2 is a 12,651 bp plasmid of the GR8/GR23 Rep3-superfamily group that encodes the *tetA(39)* tetracycline resistance determinant in a p*dif* module.

## INTRODUCTION

Infections caused by the Gram-negative pathogen, *Acinetobacter baumannii*, have become increasingly problematic, particularly among immunocompromised patients and patients in intensive care units, due to its ability to acquire and develop resistance to multiple antimicrobials thereby severely limiting treatment options (1, 2). The genomes of *Acinetobacter* strains are flexible and adaptable, prone to accumulating antibiotic resistance determinants through horizontal gene transfer involving mobile genetic elements (3, 4). Resistance to carbapenems, which are among the antimicrobials of last resort for the treatment of multidrug-resistant (MDR) *Acinetobacter* infections, is increasing with resistance rates exceeding 90% in certain regions of the world (5). Of pressing concern, pan drug-resistant (PDR) isolates of *A. baumannii*, which are resistant to all classes of antimicrobials, have been reported from clinical as well as environmental samples (6–8). In the CDC’s 2019 Antibiotic Resistance Threats Report, carbapenem-resistant *A. baumannii* has been listed as an “urgent” threat (9). Likewise, the World Health Organization (WHO) has categorized carbapenem-resistant *A. baumannii* as a critical priority pathogen towards which new antimicrobials are urgently needed (10).

*A. nosocomialis* is closely related to *A. baumannii* and along with *A. pittii, A. seifertii, A. dijkshoorniae* and *A. calcoaceticus*, they are often grouped together as the *A. baumannii-A. calcoaceticus* (Abc) complex due to difficulties in identifying these bacteria by traditional biochemical methods (11, 12). In our recent study of *Acinetobacter* isolates obtained from the main tertiary hospital in the state of Terengganu, Malaysia in 2015, the majority (83.7%) were *A. baumannii* followed by *A. nosocomialis* (10.4%) with multidrug resistance much more prevalent in *A. baumannii* (13). Nevertheless, *A. nosocomialis* and other members of the Abc complex are clinically relevant with carbapenem-resistant and MDR isolates being reported (14).

Multiple mechanisms of drug resistance are usually at play in *Acinetobacter* isolates and these include enzymatic inactivation of the antibiotic, modifications in the target sites, reduced accumulation of antibiotics through expression of efflux systems or mutations in outer membrane channels, and the formation of biofilms (15, 16). Carbapenem resistance in *Acinetobacter* is frequently attributed to the acquisition and production of OXA β-lactamases, which are categorized as Ambler class D enzymes that catalyzes the hydrolysis of the β-lactam substrate forming an intermediate covalent acyl-enzyme complex with a serine residue within the active site (17). Common acquired OXA subtypes found in *Acinetobacter* include OXA-23, OXA-24/40, OXA-58, OXA-143 and OXA-235 with the genes encoding them usually associated with or located in mobile genetic elements (16, 17). In some instances, an upstream and adjacent insertion sequence (IS) element provided a strong outward-directing promoter for their expression (17). *A. baumannii* also harbors an intrinsic *bla*_OXA-51_/*bla*_OXA-51-like_ gene in its chromosome and although the OXA-51/OXA-51-like enzyme has been shown to hydrolyze imipenem and meropenem, its affinity for these carbapenems is quite low and would not normally confer carbapenem resistance (17). However, the insertion of IS elements with outward-directing promoters such as IS*Aba1* upstream of the *bla*_OXA-51_/*bla*_OXA-51-like_ gene has been shown to increase its expression leading to carbapenem resistance (18). Nevertheless, recent reports have indicated that in the absence of an acquired carbapenemase gene, the presence of IS*Aba1* or similar elements upstream and adjacent to the intrinsic *bla*_OXA-51_/*bla*_OXA-51-like_ gene does not always guarantee carbapenem resistance in these isolates (13, 19).

The metallo-β-lactamases (MBLs) or Ambler class B enzymes, especially the New Delhi metallo-β-lactamase (NDM) group, is another class of acquired carbapenemases that have been found in *Acinetobacter* spp., being first reported in *A. baumannii* from India (20) and China (21). MBLs, including NDMs, are dependent on zinc ions at the active site of the enzyme (16). The *bla*_NDM-1_ gene has since been found in many other *Acinetobacter* spp., is usually carried in the composite transposon Tn*125* or its derivatives, and is either plasmid- or chromosomally-encoded (22). NDM-1 confers resistance to all β-lactams except monobactams such as aztreonam and are not inhibited by β-lactamase inhibitors such as clavulanic acid, sulbactam, tazobactam and avibactam (16, 23, 24). Most isolates that harbor the *bla*_NDM-1_ gene are likely MDR or extensive-drug resistant (XDR) due to the association of *bla*_NDM-1_ with other resistance genes (16, 22).

Here, we report the whole genome sequences of two NDM-1-producing *Acinetobacter* clinical isolates, *A. baumannii* AC1633 and *A. nosocomialis* AC1530, obtained from the main tertiary hospital in the eastern coast state of Terengganu in Peninsular Malaysia. We show that in these two isolates, the *bla*_NDM-1_ gene is co-located with *bla*_OXA-58_ on a large ca. 170 kb plasmid along with various antimicrobial resistance genes and that the carriage of this plasmid in these two isolates likely led to their MDR status. We also show sequence evidence that this plasmid was likely derived from two plasmids that separately encoded *bla*_NDM-1_ and *bla*_OXA-58_ in a Malaysian *A. pittii* isolate via an IS*1006*-mediated recombination event.

## RESULTS AND DISCUSSION

### Background of the *A. baumannii* AC1633 and *A. nosocomialis* AC1530 clinical isolates

*A. baumannii* AC1633 and *A. nosocomialis* AC1530 are part of our collection of *Acinetobacter* spp. clinical isolates that were obtained since 2011 from Hospital Sultanah Nur Zahirah (HSNZ), the main public tertiary hospital in the state of Terengganu, Malaysia (13, 25). Whole genome sequencing was performed on a random selection of fifty isolates obtained from 2011 – 2016 (manuscript in preparation) and preliminary analyses of the genome sequences indicated two isolates that harbored the *bla*_NDM-1_ gene, i.e., AC1633 and AC1530.

*A. baumannii* AC1633 was isolated from the blood of a 60-year old female patient in the neurology intensive care unit in April 2016. The patient had hospital-acquired pneumonia with respiratory failure and eventually succumbed to septicemia 41 days after hospital admission. *A. nosocomialis* AC1530 was isolated from the blood of a 14-year old male patient in the surgical ward in April 2015. The patient was admitted for polytrauma due to a motor vehicle accident, developed hospital-acquired pneumonia complicated with right parapneumonic effusion but recovered and was discharged after 60 days. *A. baumannii* AC1633 was resistant to the carbapenems (with imipenem, meropenem and doripenem MIC values of >32 μg/ml each), cephalosporins (cefotaxime, ceftriaxone, ceftazidime, and cefepime), β-lactam/β-lactamase inhibitor combination (piperacillin/tazobactam and ampicillin/sulbactam), trimethoprim/sulfamethoxazole, ciprofloxacin and tetracycline. AC1633 also showed resistance to gentamicin but was susceptible to the other aminoglycosides tested (namely amikacin and tobramycin), as well as to levofloxacin, doxycycline and the polymyxins (polymyxin B and colistin). On the other hand, *A. nosocomialis* AC1530 was resistant to the carbapenems (with MIC values for imipenem, meropenem and doripenem >32 μg/ml each), cephalosporins (cefotaxime, ceftriaxone, ceftazidime, and cefepime), trimethoprim/sulfamethoxazole and gentamicin but susceptible to all other antibiotics tested. Thus, both AC1633 and AC1530 are categorized as MDR following the criteria proposed by the joint commission of the United States Centers for Disease Control and Prevention (CDC) and the European Centre for Disease Prevention and Control (ECDC) (26).

### Whole genome sequencing and comparative analyses of AC1633 and AC1530

Analysis of the Illumina-sequenced genomes of *A. baumannii* A1633 and *A. nosocomialis* AC1530 indicated the presence of the *bla*_NDM-1_ and *bla*_OXA-58_ carbapenemase genes. Production of the NDM-1 metallo-β-lactamase (MBL) in both isolates was validated by testing with the E-test MBL kit (BioMérieux). Further analyses of the assembled genome data of AC1530 and AC1633 revealed the possibility that the *bla*_NDM-1_ and *bla*_OXA-58_ genes could be harbored in either one or two large plasmids in both isolates but this was difficult to ascertain as there were more than 20 assembled contigs from each isolate’s genome data that could potentially belong to these plasmids. Thus, the genomic DNA of these two isolates were subjected to PacBio sequencing and hybrid assembly was then performed on the PacBio and Illumina reads. The resulting assembled genome features of these two isolates are listed in **Table 1**.

**Table 1.** Genome features of *A. baumannii* AC1633 and *A. nosocomialis* AC1530

| Feature | *A. baumannii* AC1633 |  |  |  |  | *A. nosocomialis* AC1530 |  |
| --- | --- | --- | --- | --- | --- | --- | --- |
|  | Chromosome | pAC1633-1 | pAC1633-2 | pAC1633-3 | pAC1633-4 | Chromosome | pAC1530 |
| Size (bp) | 4,364,474 | 174,292 | 12,651 | 9,950 | 5,210 | 3,980,182 | 173,972 |
| GC content | 39% | 38% | 36% | 36% | 37% | 38% | 38% |
| Total no. of genes | 2,112 | 29 | 4 | - | 1 | 1,918 | 29 |
| No. of rRNA operons | 18 | - | - | - | - | 18 | - |
| No. of tRNAs | 72 | - | - | - | - | 74 | - |
| Total no. of coding sequences (CDS) | 4,096 | 176 | 18 | 13 | 8 | 3,750 | 180 |

The genome of *A. baumannii* AC1633 is nearly 4.4 Mb in size and is comprised of a single chromosome of 4.36 Mb (accession no. CP059300) and four plasmids designated pAC1633-1 (174 kb; CP059301), pAC1633-2 (12.6 kb; CP059303), pAC1633-3 (9.9 kb; CP059304) and pAC1633-4 (5.2 kb; CP059302). *A. baumannii* AC1633 is typed as ST2089 under the Oxford MLST scheme and ST126 under the Pasteur MLST scheme. AC1633 does not belong to any of the major *A. baumannii* global clonal lineages and phylogenetic analysis of the whole genome sequence in comparison with the sequences of selected *A. baumannii* isolates (**Fig. 1A; Suppl. Table S1**) showed that it is most closely related to *A. baumannii* CIP70.10 (ATCC 15151) which was isolated in France in 1970 and is an important reference strain due to its susceptibility to most antimicrobials (27). Average nucleotide identity (ANI) between these two isolates was determined to be 99.87%. CIP70.10 also belonged to the same STs as AC1633. Phylogenetic analyses also indicated that AC1633 is not closely related to any of the handful of Malaysian *A. baumannii* genomes that are currently available in the database, most of which belonged to the Global Clonal 2 (GC2) lineage (28, 29) except for strain PR07 (accession no. CP012035.1), which belonged to ST734 (Oxford)/ST239 (Pasteur) (30). *A. baumannii* AC12, AC29 and AC30 which were isolated from the same hospital as AC1530 and AC1633 but in the year 2011, were ST195 (Oxford)/ST2 (Pasteur) (31, 32) and showed ANI values of 97.8% in comparison with AC1633. A recent report of 13 *A. baumannii* genomes from Malaysia indicated three isolates that harbored *bla*_NDM-1_ (29) but we were unable to compare these isolates with ours as the sequence files that were associated with the GenBank accession nos. provided in the manuscript have yet to be publicly released at the time of writing (September 29, 2020). Using the KAPTIVE database which enables the typing of *A. baumannii* strains by variation in their composition and structure of capsular polysaccharide (CPS) biosynthetic genes (33), AC1633 was typed as OCL6 for the outer core biosynthesis locus and KL14 for the K locus that contained genes responsible for the biosynthesis and export of CPS, and both loci were typed with 100% match confidence level.

**Fig. 1.**
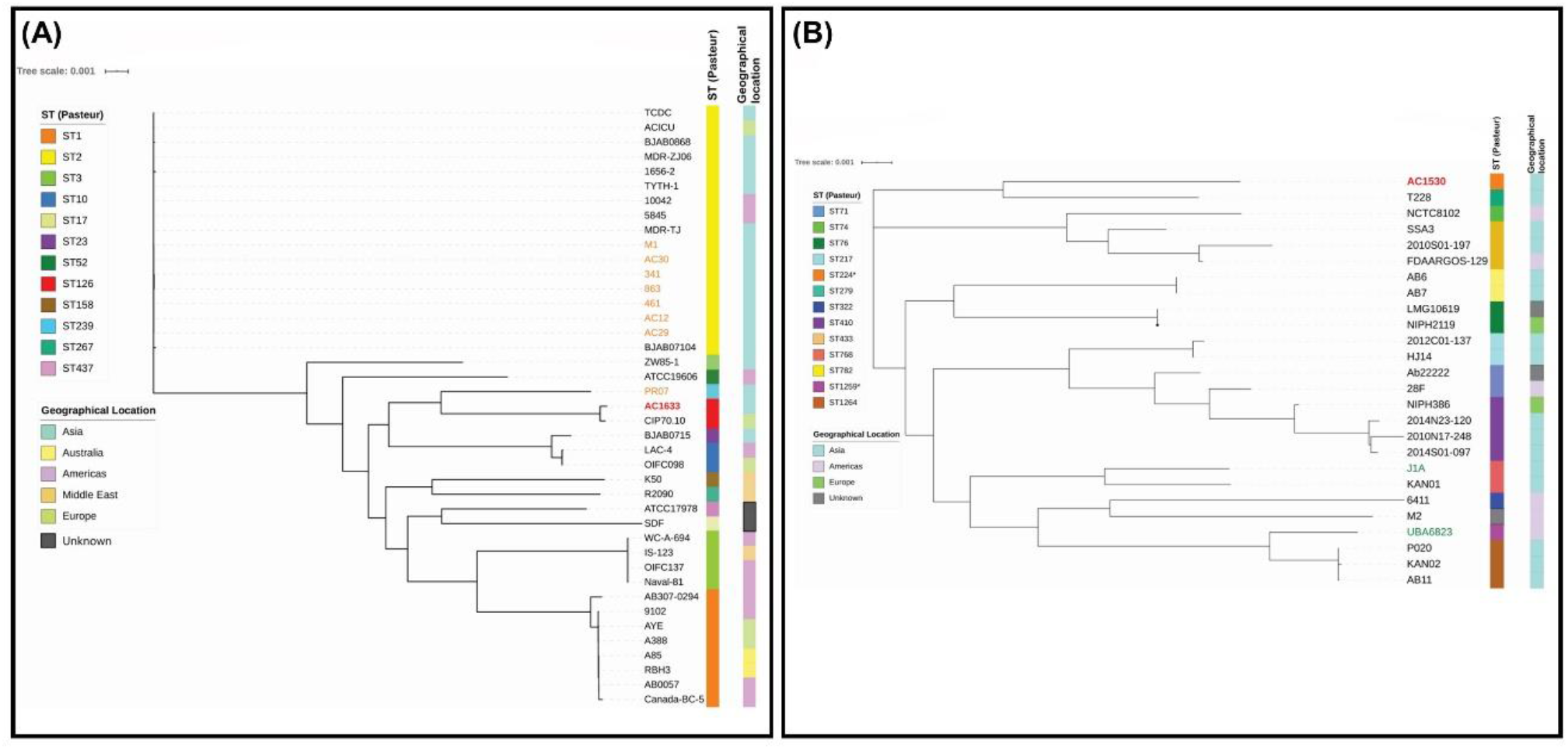
Core genome phylogenetic trees of *A. baumannii* AC1633 (A) and *A. nosocomialis* AC1530 (B) in comparison with other related isolates. The sequence types (STs) of the isolates as determined using the Pasteur scheme was presented by the colored bar on the right of the respective trees. For *A. baumannii* in **(A)**, ST1 corresponds to the Global Clone 1 (GC1) lineage, ST2 to GC2 and ST3 to GC3. The geographical location of the respective isolates was also presented as a colored bar on the furthest right of each tree. In **(A)**, *A. baumannii* isolates from Malaysia were indicated in orange fonts (except AC1633, which was indicated in bold red fonts) and in **(B)**, *A. nosocomialis* environmental isolates were indicated in blue-green fonts. All other *A. nosocomialis* isolates were obtained from clinical samples. Details of the *A. baumannii* and *A. nosocomialis* isolates that were used in the phylogenetic analyses are in **Suppl. Table 1**.

*A. nosocomialis* AC1530 has a single chromosome of 3.98 Mb (CP045560.1) and a plasmid of 173.9 kb designated pAC1530 (CP045561.1). AC1530 was assigned by the curators of the PubMLST database (34) to the Pasteur ST1539 (with alleles *cpn60-47, fusA-26, gltA-50, pyrG-14, recA-26, rplB-16* and *rpoB*-49) and Oxford ST2195 (with alleles *cpn60*-73, *gdhB*-86, *gltA*-76, *gpi*-4, *gyrB*-65, *recA*-21 and *rpoD*-90). Phylogenetic analyses showed that the closest relative of AC1530 is *A. nosocomialis* T228 (accession no. JRUA01000001.1), a clinical isolate that was obtained from Bangkok, Thailand in 2010 (**Fig. 1B; Suppl. Table S1**). However, *A. nosocomialis* T228 was typed as Pasteur ST279 and Oxford ST1897 whereby AC1530 shared only a single allele in the Oxford scheme (*gyrB*-65) and two alleles in the Pasteur scheme (*fusA*-26, *rplB*-16) with T228.

### Antimicrobial resistance genes in the genomes of AC1633 and AC1530

Interestingly, the bulk of the acquired antimicrobial resistance genes for *A. baumannii* AC1633 and *A. nosocomialis* AC1530 came from the large ca. 170 kb plasmid, pAC1633-1 and pAC1530, respectively (**Table 2**). *A. baumannii* AC1633 harbored two β-lactam resistance genes in its chromosome, i.e., the intrinsic *bla*_OXA-51-like_ gene categorized as *bla*_OXA-116_, and the *Acinetobacter*-derived AmpC cephalosporinase (ADC) gene, *bla*_ADC-25_ (accession no. EF016355.1) (35). In some cases, carbapenem resistance and increased cephalosporin resistance have been linked with upregulation of the respective *bla*_OXA-51_/*bla*_OXA-51-like_ or *blaADC* genes through insertion of IS*Aba1* or related IS elements that harbor outward-directing promoters (18, 36, 37) but no such IS elements could be found upstream of the *bla*_OXA-116_ and *bla*_ADC-25_ genes in *A. baumannii* AC1633. Tetracycline resistance in AC1633 is likely mediated by the *tetA(39)* gene that was carried in the smaller 12.6 kb plasmid, pAC1633-2, and that encode the tetracycline-specific TetA(39) efflux pump of the major facilitator superfamily (MFS). Notably, pAC1633-1 also harbors the *adeABC* operon that encodes the multidrug resistance-nodulation-cell division (RND) family efflux system along with its two-component regulatory system, *adeRS*, which is located upstream and transcribed divergently from *adeABC*. This efflux system is usually chromosomally-encoded in *Acinetobacter* and the multidrug resistance phenotype has been shown to correlate with overexpression of *adeABC* (38, 39). The chromosome of AC1633 also harbors genes encoding the other *Acinetobacter* RND family efflux pumps, *adeFGH* and *adeIJK* along with their respective regulatory genes, *adeL* and *adeN* (38, 40), three genes encoding MFS efflux pumps, i.e., *abaF, abaQ* and *amvA*, and finally, *abeS* which encode a small-multidrug resistance (SMR) family efflux pump (**Suppl. Table S2**).

**Table 2.** Antimicrobial resistance phenotype and carriage of antimicrobial resistance genes and chromosomal gene mutations in *A. baumannii* AC1633 and *A. nosocomialis* AC1530

| Strain | Antimicrobial resistance phenotype by class / carriage of resistance genes or chromosomal mutations |  |  |  |  |  |  |
| --- | --- | --- | --- | --- | --- | --- | --- |
|  | Carbapenems | Cephalosporins | Aminoglycosides | Tetracyclines | Fluoroquinolones | Sulfonamides | Macrolides |
| *A. baumannii*<br>AC1633 | IMI<sup>R</sup> MEM<sup>R</sup><br>DOR<sup>R</sup> | CTX<sup>R</sup> FOX<sup>R</sup><br>TAZ<sup>R</sup> FEP<sup>R</sup> | GEN<sup>R</sup> AMI<sup>S</sup><br>TOB<sup>S</sup> | TET<sup>R</sup> DOX<sup>S</sup> | CIP<sup>R</sup> LEV<sup>S</sup> | SXT<sup>R</sup> | ND |
| Chromosome | • *bla*<sub>OXA-116</sub><br>(*bla*<sub>OXA-51-like</sub>) | • *bla*<sub>ADC-25</sub> | - | - | • *gyrA* S81L*<br>• *parC* V104I<br>D105E* | - | - |
| pAC1633-1 | • *bla*<sub>NDM-1</sub><br>• *bla*<sub>OXA-58</sub> | - | • *aac(3)-Ild*<br>• *aph(3'')-Ib*<br>• *aph(6)-Id* | - | - | • *sul2* | • *msrE*<br>• *mphE* |
| pAC1633-2 | - | - | - | • *tetA(39)* | - | - | - |
| pAC1633-3 | - | - | - | - | - | - | - |
| pAC1633-4 | - | - | - | - | - | - | - |
| *A. nosocomialis*<br>AC1530 | IMI<sup>R</sup> MEM<sup>R</sup><br>DOR<sup>R</sup> | CTX<sup>R</sup> FOX<sup>R</sup><br>TAZ<sup>R</sup> FEP<sup>R</sup> | GEN<sup>R</sup> AMI<sup>S</sup><br>TOB<sup>S</sup> | TET<sup>S</sup> DOX<sup>S</sup> | CIP<sup>S</sup> LEV<sup>S</sup> | SXT<sup>R</sup> | ND |
| Chromosome | - | • *bla*<sub>ADC-68</sub> | - | - | • *parC* V104I<br>D105E* | - | - |
| pAC1530 | • *bla*<sub>NDM-1</sub><br>• *bla*<sub>OXA-58</sub> | - | • *aac(3)-Ild*<br>• *aph(3'')-Ib*<br>• *aph(6)-Id* | - | - | • *sul2* | • *msrE*<br>• *mphE* |

\*Mutations in the chromosomally-encoded *gyrA* and *parC* genes
*Abbreviations used:* IMI, imipenem; MEM, meropenem; DOR, doripenem; CTX, ceftriaxone; FOX, cefotaxime; TAZ, ceftazidime; FEP, cefepime; GEN, gentamicin; AMI, amikacin; TOB, tobramycin; TET, tetracycline; DOX, doxycycline; CIP, ciprofloxacin; LEV, levofloxacin; SXT, trimethoprim- sulfamethoxazole; ND, not determined; resistance, intermediate resistance and susceptibility towards the antimicrobials are denoted as superscript letters “R”, “I” and “S”, respectively.

As for *A. nosocomialis* AC1530, very few antimicrobial resistance genes are found in its chromosome, and the only resistance gene encoding an antibiotic-inactivating enzyme is a *bla*_ADC_-encoding cephalosporinase that shared 94% amino acid sequence identity with ADC-68 (accession no. AGL39360.1)(41). However, as in the *A. baumannii* AC1633 genome, no IS elements with outward-directing promoters could be detected upstream of this gene. Efflux pumps that are encoded in the AC1530 chromosome are the RND family *adeFGH* and its regulatory gene, *adeL*, the MFS superfamily pumps *amvA* and *abaQ*, and the SMR family *abeS* (**Suppl. Table S2**). The same suite of resistance genes in pAC1633-1 was found in pAC1530 (**Table 2**) and therefore, likely contribute to its resistance phenotype.

### Characteristics of pAC1633-1 and pAC1530 and their carriage of antimicrobial resistance genes

Plasmids pAC1653-1 from *A. baumannii* AC1633 and pAC1530 from *A. nosocomialis* AC1530 were nearly identical except at five regions (**Fig. 2**): (i) insertion of IS*Aba11* into an ORF encoding an 85-aa hypothetical protein (locus tag: GD578_19675; accession no. QGA46103.1) which contains a ribosomal protein L7/L12 C-terminal domain (pfam00542) in pAC1530 at nt. 40824 (nts. 40825 – 41927 of pAC1633-1). This ORF is upstream of the *traMNO* genes and lies within a cluster of genes that are proposed to be part of the conjugative transfer region for the plasmid. (ii) An I*S4* family transposase at nts. 58863 – 60133 in pAC1633-1 with no matches to existing IS elements in the ISFinder database (42) and no inverted repeats flanking the putative transposase gene. This is likely a remnant of an IS element that had inserted within IS*Aba31*, leading to only a partial IS*Aba31* downstream of this IS*4*-family remnant element. A full-length IS*Aba31* is found in the corresponding site in pAC1530 with a characteristic 2-bp “TA” direct repeat flanking the IS element. (iii) A 255 bp insertion within a hypothetical ORF at nt. 89589 of pAC1530 in pAC1633-1 (nts. 91978 – 92233). No characteristic signature sequences of mobile elements could be detected within this short fragment. (iv) Insertion of IS*Aba11* into an ORF encoding a putative toxin of the Zeta toxin family in pAC1530. This insertion, at nt. 101270 of pAC1633-1, led to a 5 bp direct repeat (“TATAG”) in pAC1530 (nts. 98631 – 99731). (v) Addition of a *relBE* toxin-antitoxin (TA) system along with a downstream ORF encoding a protein of the SMI1/KNR4 family in pAC1530 at nt. 164446 of pAC1633-1. This 1,192 bp fragment is flanked by *pdif* (XerD/XerC) recombination sites, of which will be covered in more detail in a later section. pAC1633-1 and pAC1530 belong to a group of diverse *Acinetobacter* plasmids that do not have an identifiable replication initiator or replicase (Rep) protein (43, 44).

**Fig. 2.**
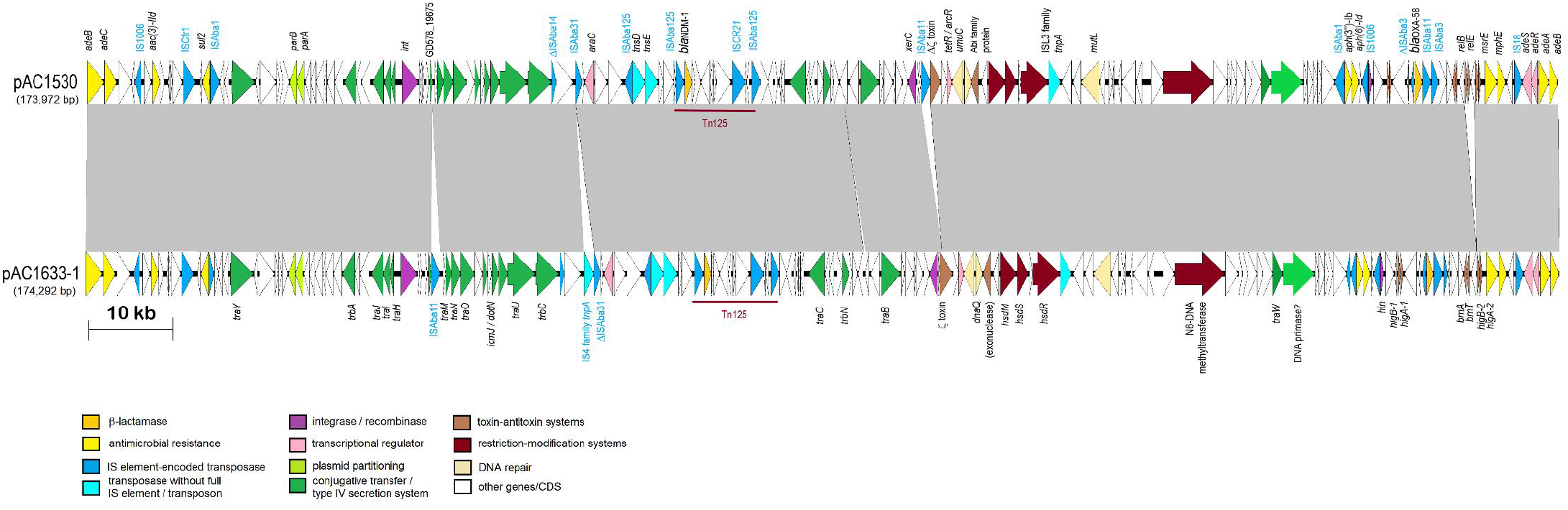
Comparative linear map of plasmids pAC1633-1 and pAC1530. Arrows indicate the extents and directions of genes and ORFs. The *bla*_NDM-1_ and *bla*_OXA-58_ genes are colored in gold, other antimicrobial resistance genes are in yellow. Putative transcriptional activators are in pink, including the genes encoding the two-component regulatory proteins, *adeR* and *adeS* that controls transcription of the *adeABC* efflux pump. Transposases encoded by full copy IS elements are shown in dark blue whereas transposases without their corresponding IS elements or transposons in full are depicted in light blue. Genes with homologies to conjugative transfer or type IV secretion system genes are in dark green. GD578_19675 refers to the ORF in the conjugative region of pAC1530 that was the site of insertion for IS*Aba11* in pAC1633-1. Color codes for the other genes are as indicated. Tn*125* that harbors the *bla*_NDM-1_ gene is labeled. The extent of regions with >99% nucleotide sequence identities are indicated in the grey-shaded area.

Both pAC1633-1 and pAC1530 are a repository of several antimicrobial resistance genes including the *bla*_NDM-1_ and *bla*_OXA-58_ carbapenemase genes (**Table 2**). Three aminoglycoside resistance genes were found in both plasmids and all three encoded for aminoglycoside-modifying enzymes. The *aac(3)-IId* is a subclass of the AAC(3) enzymes that catalyze acetylation of the -NH_2_ group at the 3-position of the aminoglycoside’s 2-deoxystreptamine nucleus and usually confers resistance to gentamicin, sisomicin and fortimicin (45). Both *A. baumannii* AC1633 and *A. nosocomialis* AC1530 were gentamicin resistant but their susceptibilities against sisomicin and fortimicin were not tested. The other two aminoglycoside resistance genes found in pAC1633-1 and pAC1530 encode the aminoglycoside O-phophotransferases (APHs), *aph(6)-Id* and *aph(3”)-Ib*, and both genes confer resistance to streptomycin (which was, however, not phenotypically tested in both *Acinetobacter* strains). Both genes were adjacent to each other and are flanked by IS elements with IS*Aba1* upstream of *aph(3”)-Ib* and IS*1006* downstream of *aph(6)-Id* (**Fig. 2**). The contiguity of *aph(3”)-Ib* and *aph(6)-Id* was initially reported in the broad-host range IncQ plasmid RSF1010 where they were part of a fragment that included the genes *repA, repC, sul2, aph(3”)-Ib* and *aph(6)-Id* that has later been found, complete or in part, within plasmids, integrative conjugative elements and genomic islands (45, 46).

The *sul2* gene that encodes for dihydropteroate synthase which confers sulfonamide resistance is sandwiched between IS*Aba1* upstream and IS*Cfr1* downstream in both pAC1633-1 and pAC1530 (**Fig. 2**). In the chromosome of *A. baumannii* ATCC 19606, *sul2* is associated with the IS*CR2* element and is part of a large (36,157 bp) genomic island designated GI*sul2* (47). The association of *sul2* with the IS*CR2* element was previously reported in the plasmid RSF1010 (48). However, in the GC1 *A. baumannii* RUH875, IS*Aba1* was detected upstream of *sul2* and provided a promoter for its expression (49). This was observed in pAC1633-1 and pAC1530 but in these two plasmids, the IS*CR2* element was truncated due to the insertion of the 1,617 bp IS*Cfr1*. No direct repeats were found at the site of IS*Cfr1* insertion, as was reported for this IS element in ISFinder (https://isfinder.biotoul.fr/scripts/ficheIS.php?name=ISCfr1).

The three aminoglycoside resistance genes, *aac(3)-IId, aph(6)-Id* and *aph(3”)-Ib* along with the *sul2* gene were found to be in a 42,125 bp fragment in pAC1530 that was flanked by IS*Aba1* with characteristic 9-bp direct repeat of the target sequence in a typical composite transposon-like structure. This 42 kb fragment also included the *bla*_OXA-58_, *msrE, mphE, adeRS-adeABC* resistance genes that were nested in a 29,670 bp fragment flanked by IS*1006* (**Suppl. Fig. 1**) and which is postulated to be derived from a smaller, separate plasmid via IS*1006*-mediated recombination/transposition. Comparative sequence analysis subsequently indicated that the IS*Aba1*-flanked composite transposon, designated Tn*6948* by the Transposon Registry (50), is 14,750 bp and details of its structure and how the 42 kb region in pAC1530 (as well as pAC1633-1) came about will be presented in a later section of this manuscript.

The *bla*_NDM-1_ gene is found within a 10,099 bp Tn*125* composite transposon that was made up of a pair of flanking IS*Aba125* elements and is a common genetic vehicle for the dissemination of *bla*_NDM_ genes in *Acinetobacter* spp. (22, 23, 51). One copy of the IS*Aba125* is 93 bp upstream of *bla*_NDM-1_ and the presence of an outward-directing promoter [a typical σ^70^-type promoter with the –35 sequence (TTGAAA) separated by 16 bp to the –10 sequence (TTGAAT)] at the terminal inverted repeat of IS*Aba125* likely drives the expression of *bla*_NDM-1_ (23, 52). In both pAC1633-1 and pAC1530, *Tn125* was inserted into an unknown open reading frame resulting in a 3 bp target site duplication (“ACG”) (**Fig. 3**), as has been previously reported for this transposon (23, 51). However, a recent publication of a 265 kb plasmid pABF9692 that also co-harbored *bla*_NDM-1_ and *bla*_OXA-58_ from a pandrug-resistant *A. baumannii* ABF9692 chicken isolate revealed a 4 bp duplication (“CCAT”) at the site of insertion of Tn*125* (53). The genome of *A. baumannii* JH was reported to harbor Tn*125* bracketed by 3-bp target site (“TTC”) duplication (51). However, upon closer inspection of the DNA sequence (accession no. JN872329), we found that the target site duplication was 4-bp (“TTCC”) instead of 3-bp (nts. 159 – 162 on the left flank and nts. 10262 – 10265 on the right flank of the transposon). Although experimental evidence for the transposition of Tn*125* had revealed that its insertion always led to 3-bp duplication of the target sequence (52), there are thus instances in natural isolates whereby 4-bp target site duplication were observed.

**Fig. 3.**
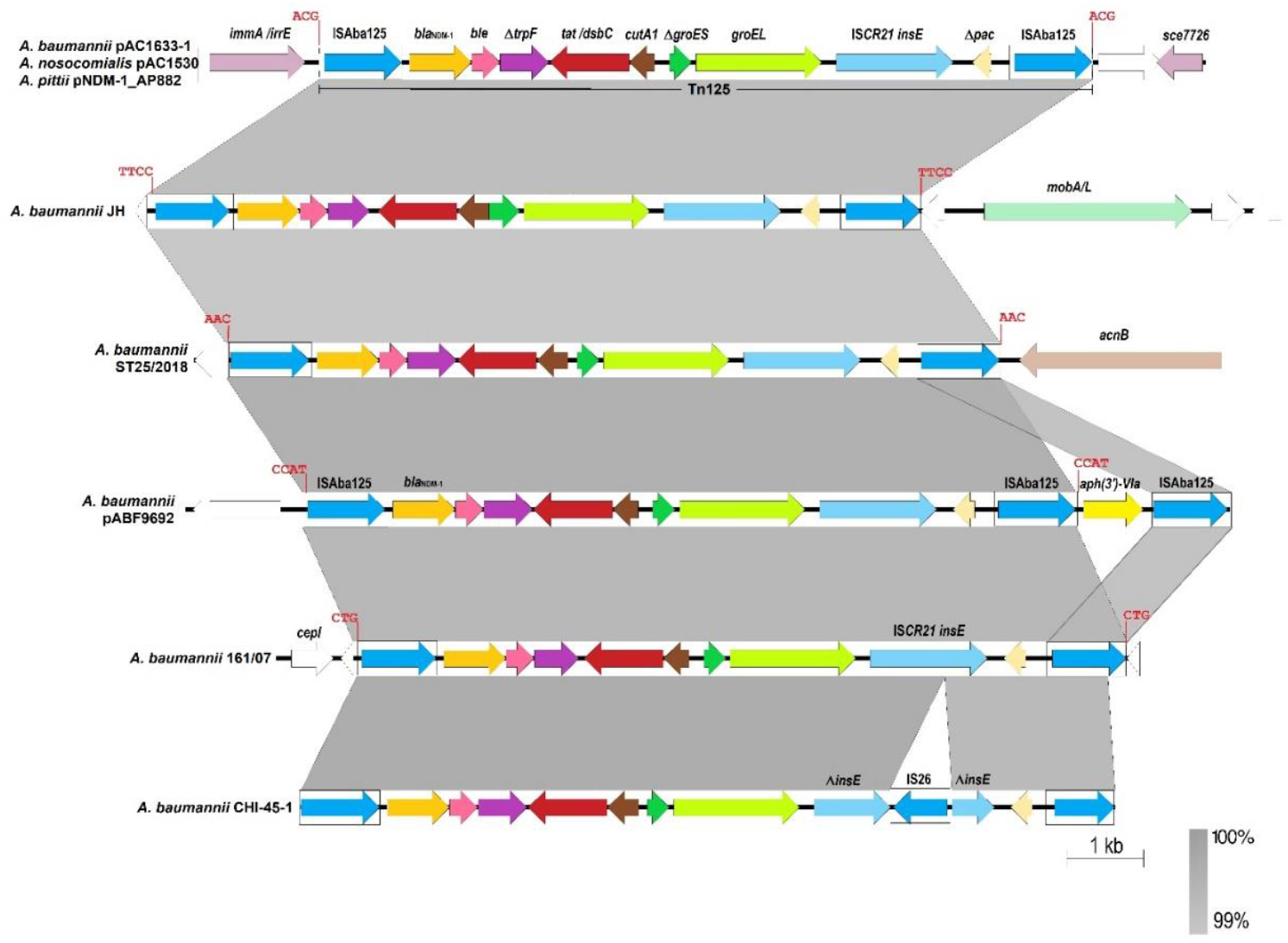
Comparative map of Tn*125* and its surrounding genes in several *Acinetobacter* isolates. Arrows indicate the extents and directions of the genes and ORFs with the *bla*_NDM-1_ gene shown in gold color. IS elements are indicated in boxes with their encoded transposases depicted as blue arrows. The transposase encoded by the *ISCR21* element, which is capable of mobilizing genes at its left-hand extremity by rolling circle transposition (51), is shown in light blue arrow and labeled as *“insE”*. The Tn*125* target site duplications identified in each isolate are shown in bold red fonts above their site of insertions. Note that for *A. baumannii* CHI-45-1 (accession no. KF702386), no target site duplication could be identified as the GenBank entry contained only the Tn*125* sequence and not its surrounding sequences. Genes within Tn*125* are as follows: *ble*, gene conferring resistance to bleomycin; Δ*trpF*, truncated phophoribosylanthranilate isomerase (sometimes designated as *iso*); *tat/dsbC*, twin-arginine translocation pathway signal sequence protein; *cutA1*, divalent cation tolerance protein (also designated *dct*); *groES* and *groEL*, chaperonin proteins; Δ*pac*, truncated phospholipid acetyltransferase. In pAC1633-1, pAC1530 and pNDM-1_AP882, the *immA/irrE* gene encodes for a protein of the ImmA/IrrE family metalo-endopeptidase and *sce7726* encodes for a protein of the sce7726 family; in *A. baumannii* JH (accession no. JN872329.1), *mobA/L* encodes a mobilization protein of the MobA/L family; in *A. baumannii* ST25/2018 (accession no. MK467522.1), *acnB* encodes aconitate hydratase B; and in *A. baumannii* 161/07 (accession no. HQ857107), *cepI* encodes homoserine lactone synthase while Tn*125* was inserted into the *mfs* gene that encodes a transport protein of the major facilitator superfamily (96). White arrows refer to ORFs that encode hypothetical proteins. For *A. baumannii* plasmid pABF9692 (accession no. CP048828), nts 191,490 – 205,120 was covered in the comparative analysis, whereas for the *A. pittii* AP882 pNDM-1_AP882 plasmid (accession no. CP014478), the analysis covered nts. 67,185 – 80,160. In the case of pAC1633-1, the depicted map covered nts. 70,519 – 83,520, and for pAC1530, the coverage was from nts. 68,131 to 81,120. The extent of regions with 99 – 100% nucleotide sequence identities are shown in grey.

### The *bla*_OXA-58_, the *msrE-mphE* resistance genes and toxin-antitoxin systems are in p*dif* modules in pAC1633-1 and pAC1530

One of the intriguing features of *Acinetobacter* plasmids is the presence of discrete modules flanked by conserved inverted repeats homologous to the XerC and XerD binding sites (*dif* sites) separated by a 6 bp spacer, which are recombination targets for the XerC and XerD proteins (54–56). Since their initial discovery flanking a discrete module encoding the OXA-24 carbapenemase gene in the *A. baumannii* pABVA01a plasmid (57), several of these designated p*dif* modules (named for plasmid *dif*) which comprise of a pair of inverted p*dif* sites surrounding a gene or several genes, have been described harboring antimicrobial and metal resistance genes, toxin-antitoxin systems as well as other genes (56, 58). In pAC1633-1 and pAC1530, the region surrounding *bla*_OXA-58_ is rich in p*dif* sites: 11 of the 14 p*dif* sites in pAC1530 were located within a 14,410 bp fragment that spanned nts. 153,566 to 167,912 (**Fig. 4**).

**Fig. 4.**
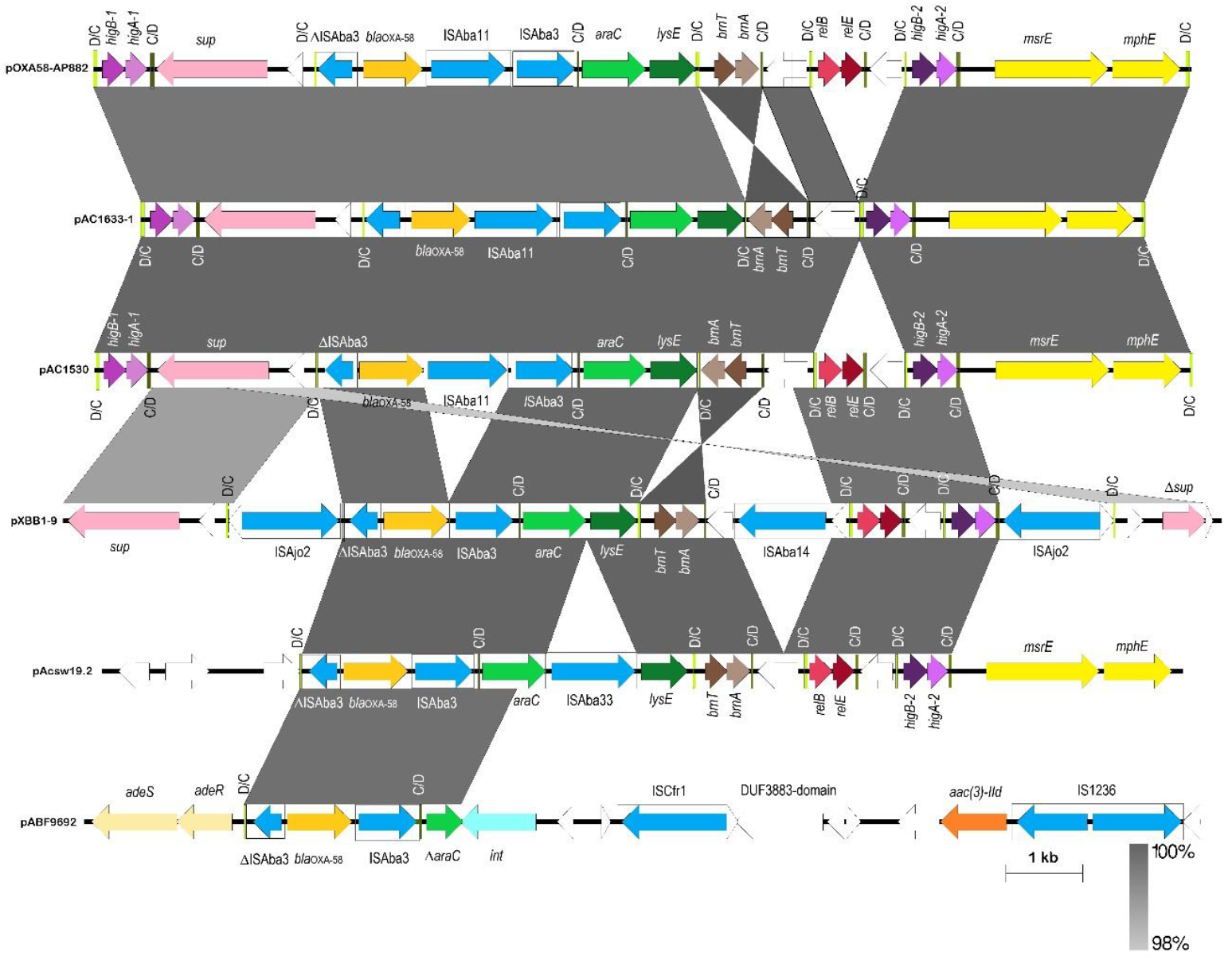
Comparative map of the p*dif*-rich regions surrounding the *bla*_OXA-58_ gene in several *Acinetobacter* plasmids. Arrows indicate the extents and directions of the genes and ORFs with the *bla*_OXA-58_ gene depicted as a gold arrow, the *msrE* and *mphE* macrolide resistance genes are in yellow and the aminoglycoside resistance gene *aac(3)-IId* is shown in orange. IS elements are shown as boxes with their encoded transposases in blue arrows within their respective boxes. p*dif* sites are depicted as vertical bars with the orientation of the sites labeled and colored as follows: XerD/XerC colored lime-green and labeled as “D/C”, XerC/XerD colored dark olive-green and labeled as “C/D”. Note the toxin-antitoxin genes that make up the following p*dif* modules: *higBA-1, higBA-2, brnTA* and *relBE*. Other genes are labeled as follows: *araC*, putative transcriptional regulator of the AraC family; *lysE*, putative threonine efflux protein; *sup*, putative sulfate transporter. White arrows depict ORFs that encode hypothetical proteins. Accession numbers and coverage of plasmid regions for the comparative map are as follows: pOXA-58_AP882 (accession no. CP014479; nts. 25,121 – 36,862 and continued from 1 – 2,560); pAC1633-1 (accession no. CP059301; nts. 155,106 – 168,160); pAC1530 (accession no. CP045561.1; nts. 153,566 – 167,840); pXBB1-9 (accession no. CP010351; nts. 2,158 – 1 and continued from nts. 398,857 – 398,921); pAcsw19.2 (accession no. CP043309; nts. 47,521 – 61,600); and pABF9692 (accession no. CP048828; nts. 113,261 – 127,790). The extent of regions with nucleotide sequence identities of between 98 – 100% are shown in grey.

The *bla*_OXA-58_ gene itself is flanked by IS elements (a partial 427-bp IS*Aba3* upstream of *bla*_OXA-58_ and full copies of IS*Aba11* and IS*Aba3* immediately downstream), which are in turn, flanked by a pair of inverted p*dif* sites, an arrangement that has been previously reported for the *A. johnsonnii*-encoded plasmid pXBB1-9, but without the presence of IS*Aba11* (59). In pAC1633-1/pAC1530, insertion of IS*Aba11* led to a characteristic 5 bp direct repeat (“ATTTA”) of the target sequence. In some *Acinetobacter*, a full IS*Aba3* or IS*Aba3*-like element is found upstream of *bla*_OXA-58_ but in other instances, this upstream IS*Aba3* is disrupted by other IS elements (55, 60–62).

A pair of inversely oriented p*dif* sites was also found to flank the *msrE* and *mphE* macrolide resistance genes in pAC1633-1 and pAC1530 (**Fig. 4**) leading to the formation of a 2,950 bp macrolide resistance p*dif* module, as had been reported previously in other *Acinetobacter* plasmids (55). This *msrE-mphE* module was always found adjacent to a *higBA* TA p*dif* module (55) and this was also the case in pAC1633-1 and pAC1530.

Intriguingly, all known TA systems detected in pAC1633-1 and pAC1530 were found within this region and they were each flanked by a pair of inverted p*dif* sites. Thus, the *higBA-1, brnTA* and *higBA-2* TA systems that in both plasmids qualify as bona-fide p*dif* modules. As mentioned earlier, one of the differences between pAC1633-1 and pAC1530 is the addition of a *relBE* TA system and an ORF encoding a protein of the SMI1/KNR4 family in pAC1530 upstream of *higBA-2* (**Fig. 4**). The *relBE* genes and the SMI1/KNR4 protein-encoded gene are each p*dif* modules as they are flanked by a pair of inverted p*dif* sites. A closer examination of the p*dif* (XerD/C) sequences showed that in pAC1633-1, the p*dif* (XerD/C) sequences upstream of *higBA-2* is a hybrid of the p*dif* (XerD/C) sequences that flanked the *relBE*-SMI1/KNR4 module in pAC1633-1: the sequences of the XerD site are identical to the sequences found upstream of *relBE* while the 6-bp spacer and the XerC site sequences are identical to the sequences upstream of *higBA-2* in pAC1633-1 (**Suppl. Fig. 2**). This suggests that the *relBE*-SMI1/KNR4 p*dif* module could have been deleted from pAC1530 in pAC1633-1 via a Xer-mediated recombination event.

Another Xer-related rearrangement could be seen when comparing the sequences of pAC1633-1 and pAC1530 with their closest plasmid relative, pOXA-58_AP882 whereby the *brnTA* TA p*dif* module is found to be in inverted orientation. The orientation of *brnTA* in pOXA-58_AP882 is, however, the same in pXBB1-9 and pAcsw19.2 (**Fig. 4**). Interestingly, the XerD and the 6-bp spacer sequences of the flanking p*dif* modules for *brnTA* are identical when comparing pAC1633-1/pAC1530 with pOXA-58_AP882 but the XerC sequences are inverted (pAC1633-1/pAC1530: TTATGCGAAGT; pOXA-58_AP882: ACTTCGCATAA) (**Suppl. Fig. 2**). Currently, genome sequencing data strongly supports the likelihood that p*dif* modules are mobile although to our knowledge, there has yet to be any definite experimental evidence offered or mechanism of mobility elucidated (54). Deletions and inversions of *Acinetobacter* p*dif* modules were hinted at (54, 58) and here, we show sequencing evidence that these do occur.

Both pAC1633-1 and pAC1530 encode for a 298 aa-residue recombinase-like protein (nts. 100,682 – 99,786 in pAC1633-1; nts. 98,036 – 97,140 in pAC1530) that was annotated as “tyrosine-type recombinase/integrase” by the NCBI Prokaryotic Genome Annotation Pipeline (PGAP) but was annotated as “*xerC*” by PROKKA. The protein encoded by this ORF shares 39% amino acid sequence identity with the corresponding chromosomally-encoded XerC and 32% identity with XerD. This *xerC*-like gene is itself flanked by p*dif* modules with the XerC/D site upstream of the gene being less identical to the other XerC/D sequences in the plasmid (3/11 nucleotide differences in the consensus XerC site and 4/11 nucleotide differences in the consensus XerD site). So far, only the 398.9 kb pXBB1-9 of *A. johnsonnii* was reported to encode both *xerC* and *xerD* within the plasmid and clustering of the p*dif* sites were also reported around the *bla*_OXA-58_ region of this plasmid (59). Whether the product of this plasmid-encoded *xerC*-like gene or the chromosomally-encoded XerCD are involved in the mobility of the p*dif* modules will require future experimental validation.

The plasmid-encoded *xerC*-like gene is divergently transcribed from a gene encoding a Zeta-like toxin which is interrupted by ISAba11 in pAC1530. This putative Zeta-like toxin, at 501 aa residues, is much larger than canonical Zeta toxins (~270 aa) of the Epsilon-Zeta/PezAT TA systems (63, 64) and is even larger than the 360 aa Zeta-like toxins of several *Acinetobacter* plasmids that had been previously characterized as “non-functional” toxins (65). There was conservation of amino acid residues within the Walker A motif of Zeta toxins which function to bind ATP for phosphorylation reactions but no conservation of amino acids that bind to the substrate for Zeta, UDP-N-acetylglucosamine (UNAG) (63) was observed for the pAC1633-1-encoded Zeta. This infers that the pAC1633-1-encoded Zeta is probably a functional kinase but has a different substrate to Zeta/PezT toxins. The absence of a candidate Epsilon-like antitoxin adjacent to this Zeta-like toxin gene also suggests that the pAC1633-1-encoded Zeta is most probably not part of a bona fide TA system and has a different function to that of known Zeta toxins.

### pAC1633-1/pAC1530 is likely a hybrid of two plasmids with the co-integration mediated by IS*1006*

BLASTN analysis of pAC1633-1 and pAC1530 showed that they are most similar in sequence to two plasmids found in *Acinetobacter pittii* AP882 designated pNDM-1_AP882 (accession no. CP014478) and pOXA-58_AP882 (accession no. CP014479). Notably, *A. pittii* AP882 was isolated from Peninsular Malaysia in 2014 but from a different state (Perak) (66) as compared to AC1633-1 and AC1530 (isolated from Terengganu in 2016 and 2015, respectively). When comparing the 146,597 bp pNDM-1_AP882 with pAC1530 and pAC1633-1, pNDM-1_AP882 is nearly identical with pAC1530/pAC1633-1 except for a 1,940 bp region adjacent to IS*1006*. This region contains a 1,472 bp IS-like sequence (nts. 4,480 – 1,472) that encodes two transposases characteristic of the IS*3* family flanked by 21/22 bp imperfect inverted repeats and 5 bp direct repeat (“ACCTG”) of the target sequence (**Suppl. Fig 3**). Further analysis of pNDM-1_AP882 led to the discovery of a 14,750 bp composite transposon designated Tn*6948* formed by flanking IS*Aba1* sequences with 9-bp target site duplication (“TTAAAAATT”) that is characteristic of this IS element (**Suppl. Fig 1**). The target site duplication is only found for the entire transposon structure but not for each individual IS*Aba1* element, thus inferring that this is likely an active transposon. Tn*6948* harbors the *sul2* sulfonamide resistance gene and the aminoglycoside resistance genes *aph(3”)-Ib, aph(6)-Id* and *aac(3)-IId. Tn6948* has an overall GC content of 50.6% as compared to 38-39% of the surrounding genes, thus suggesting its possible non-*Acinetobacter* origin.

When comparing the 36,862 bp pOXA-58_AP882 plasmid with pAC1530 and pAC1633-1, a 29,671 bp fragment of pOXA-58_AP882 was found to be identical to pAC1530/pAC1633-1 and this region, which comprises the resistance genes *bla*_OXA-58_, *msrE, mphE*, and *adeABC-adeRS*, is flanked by two copies of IS*1006* (**Fig. 5**). The remaining 7,191 bp of pOXA-58_AP882 that was absent in pAC1530/pAC1633-1 encodes 10 ORFs and this includes ORFs that encode a MobA/L mobilization protein, a Rep3 family *Acinetobacter* replicase of the GR12 group (67), a hypothetical protein with a helix-turn-helix motif that had previously been misannotated as RepA (44), a protein of the RelE/ParE toxin family and downstream of it, an ORF that encodes another helix-turn-helix protein but of the Xre family (**Suppl. Fig. 4**). The putative RelE/ParE family toxin in this region of pOXA-58_AP882 shared only 27% amino acid sequence identity with the RelE toxin of the previous RelBE TA pair found within this plasmid as well as in pAC1530 (but which was absent in pAC1633-1). In comparing the IS*1006* sequences in these plasmids, it was found that the two IS*1006* copies in pOXA-58_AP882 were identical with the two copies in pAC1530 and pAC1633-1. However, the solitary IS*1006* copy in pNDM-1_AP882 had a single nucleotide change in which T replaced C at nt. 175 of the 819 bp IS*1006*.

**Fig. 5.**
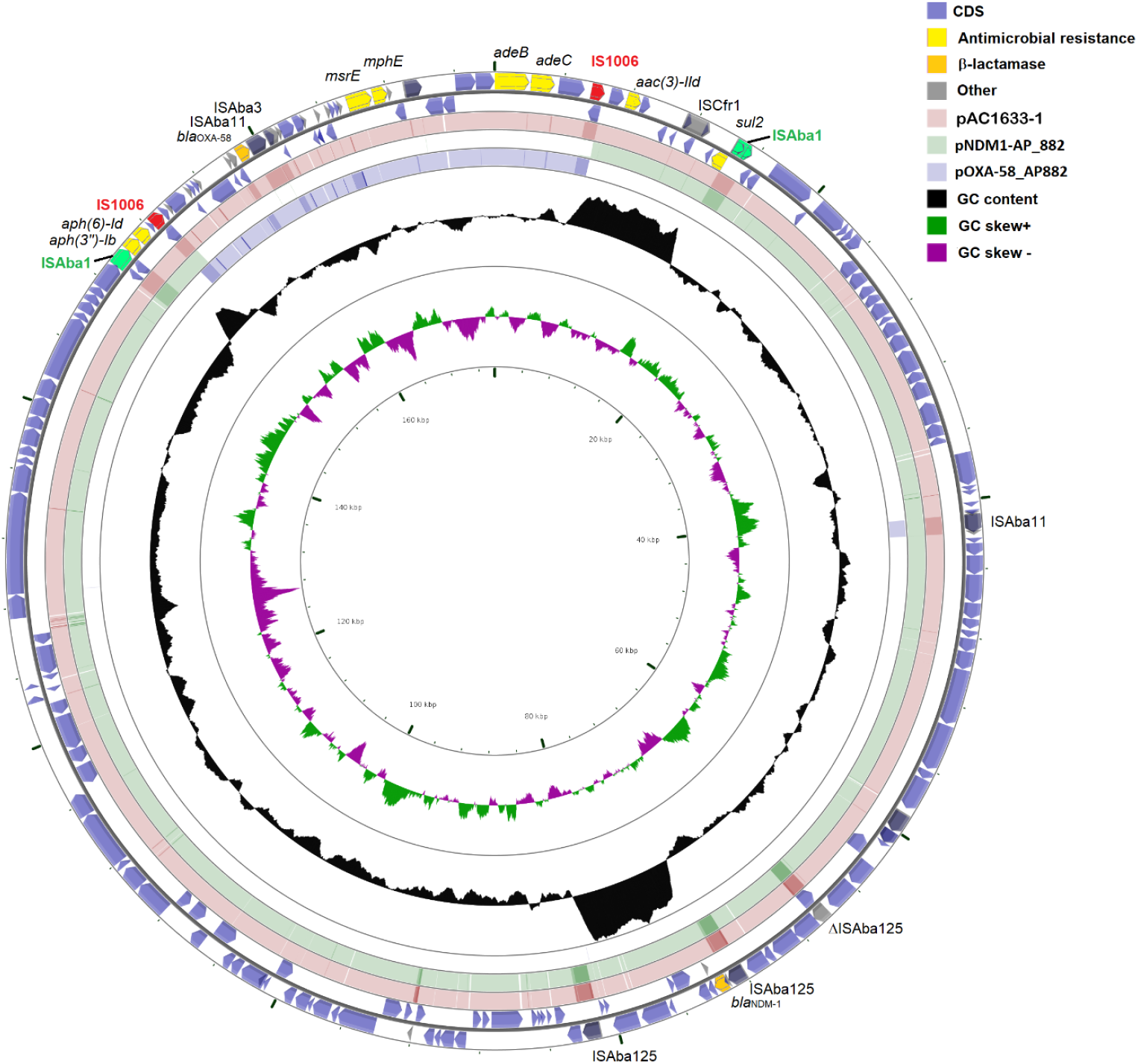
Comparison of pAC1633-1 with plasmids pNDM-1_AP882 and pOXA-58_AP882 from *Acinetobacter pittii* AP882. The outer two circles show the genes and coding sequences (CDS) from pAC1633-1 with the two copies of IS*1006* marked in red and the antimicrobial resistance genes in yellow. The two IS*Aba1* elements that flank the composite transposon Tn*6948* in pNDM-1_AP882 are shown in green. The pink-colored ring indicates pAC1633-1 while the green- and blue-colored inner rings show regions of pNDM-1_AP882 and pOXA-58_AP882, respectively, that shared >95% nucleotide sequence identities with the corresponding region in pAC1633-1. Darker shades of pink, green and blue indicate repeat regions (usually IS elements). Note how pAC1633-1 could possibly came about through integration of the IS*1006*-flanked region which encompassed the *bla*_OXA-58_, *msrE* and *mphE* resistance genes into the single IS*1006* copy of pNDM-1_AP882, as illustrated in **Suppl. Fig. 5.**

It is thus tempting to speculate that the 29,671 bp region of pOXA-58_AP882 which contained *bla*_OXA-58_ formed a composite transposon-like structure flanked by two copies of IS*1006* and this region could have transposed or recombined with pNDM-1_AP882 at its single IS*1006* copy that resided within Tn*6948* resulting in a predecessor for plasmids pAC1530 and pAC1633-1 which contained both the *bla*_NDM-1_ and *bla*_OXA-58_ genes in a single plasmid that has two copies of IS*1006* (**Suppl. Fig. 5**). IS*1006* belongs to the large IS*6*/IS*26* family of IS elements (68) and this family, in particular IS*26*, has been known to mediate the formation of cointegrates between two DNA molecules with the donor molecule harboring IS*26* (69). However, this route, designated “replicative” or “copy-in” usually leads to the formation of 8-bp target site duplication for the IS*26* and inserts at random sites (68). Here, no target site duplication could be detected in pAC1530, pAC1633-1 or even pOXA-58_AP882 at the ends of the IS*1006*-flanked region. Analysis of all the individual IS*1006* copies in pAC1530, pAC1633-1, pOXA-58_AP882 and pNDM-1_AP882 showed no evidence of target site duplications flanking each IS*1006* copy and no target site duplications were also recorded for the IS*1006* entry in the ISFinder database (https://isfinder.biotoul.fr/scripts/ficheIS.php?name=IS1006). Nevertheless, IS*26* was recently demonstrated to perform a unique transposase-dependent reaction when both donor and target molecules carry a copy of IS*26*. This reaction, designated “targeted conservative”, is targeted, occurring at one or the other end of the two IS*26* elements and with the IS element not duplicated and a target site duplication not generated (70). Cointegration by the targeted conservative route was found to be the preferred reaction if two copies of IS*26* in two different DNA molecules are available (70, 71). Based on sequence analysis alone, it is difficult to ascertain the mechanism by which the predecessor cointegrate plasmid for pAC1530 and pAC1633-1 was formed – whether it is through the “targeted conservative” route since both pOXA-58_AP882 and pNDM-1_AP882 harbored IS*1006*, or by homologous recombination via IS*1006*, or even by classical transposition as the two copies of IS*1006* that flanked the 29,671 bp *bla*_OXA-58_ fragment do form a composite transposon structure albeit without the characteristic target site duplications at its termini.

### Transmissibility of pAC1530 and pAC1633-1

The fact that pAC1530 and pAC1633-1 were nearly identical, large (>170 kb) plasmids that were isolated from two different *Acinetobacter* species in two different years (*A. nosocomialis* AC1530 from 2015 and *A. baumannii* AC1633 from 2016) but from the same hospital is suggestive of plasmid transmissibility. Sequence analysis also indicated the presence of several conjugative transfer-related genes, most of which shares between 50 – 70% amino acid sequence identities with the corresponding translated proteins of the conjugative plasmid pA297-3 from *A. baumannii* A297 (72) (**Table 3**). The conjugative transfer genes of pAC1530 and pAC1633-1 were broadly distributed in two large regions of the plasmids, as were in pA297-3. The order of the transfer genes in both regions in pAC1530 and pAC1633-1 (designated Regions 1 and 2) was identical with that in pA297-3 even though their nucleotide sequence identities were lower than 65% in some parts of these two regions (**Fig. 6**). However, in both pAC1530 and pAC1633-1, Region 1 which spans from *traW* to *trbC*, was interrupted by a 42 kb fragment encompassing the IS*Aba1*-flanked composite transposon Tn*6948*, and nested within it, the 29 kb IS*1006*-flanked fragment derived from pOXA-58_AP882 and which encode resistance genes such as *bla*_OXA-58_, *mphE-msrE*, and *sul2* (**Fig. 6**). Conjugation assays were performed using the carbapenem resistant parental hosts, *A. baumannii* AC1633 and *A. nosocomialis* AC1530, as donor strains and carbapenem susceptible *A. baumannii* ATCC 19606 and *A. baumannii* AC1529 clinical isolate that were induced to sodium azide resistance with MIC values of >300 μg/ml as recipients. Despite repeated attempts with established conjugation assay protocols (72, 73) and using different ratios of donor to recipient cells, no transconjugants were obtained that were able to grow on the selection plates (LB agar supplemented with 10 μg/ml imipenem and 300 μg/ml sodium azide). Thus, we were unable to provide direct laboratory experimental evidence that pAC1530 and pAC1633-1 were transmissible. The 200 kb pA297-3 plasmid from *A. baumannii* A297 which encoded the *sul2* sulfonamide and *strAB* streptomycin resistance genes, was found to transfer sulfonamide and streptomycin resistance to a rifampicin-resistant *A. baumannii* ATCC 17974 strain at a high frequency of 7.20 × 10^-2^ transconjugants/donor (72). However, two other plasmids, pD4 and pD46-4, which shared the transfer regions with pA297-3 were found to be non-transmissible (74, 75). In the case of pD4, an IS*Aba25*-like element was inserted into the DNA primase gene downstream of *traW*, indicating the possibility that this gene could be involved in conjugative transfer (74). However, for pD46-4, no such IS or other genetic elements were found to have interrupted the conjugative transfer-related genes; besides, no SNPs were detected in the transfer genes that might have led to have led to a frameshift or a premature stop codon within these genes (75). The reason for the apparent non-transmissibility of pD46-4 as compared to pA297-3 was not known (75). As for pAC1530 and pAC1633-1, their apparent non-transmissibility could be attributed to the insertion of the 42 kb fragment containing Tn*6948* and the IS*1006*-flanked resistance region from pOXA-58_AP882 into the conjugative transfer region 1. However, the insertion of Tn*6948* at the same site was already apparent in pNDM-1_AP882 although here, the insertion was only 14.2 kb (**Suppl. Figs. 1 and 3**). Since we do not have access to the *A. pittii* AP882 strain that harbored pNDM-1_AP882 as well as pOXA-58_AP882, we were thus unable to experimentally determine if pNDM-1_AP882 is transmissible. Nevertheless, the genomic sequence evidence presented here strongly infers the transmissible nature of pNDM-1_AP882 and by extension, pAC1530 and pAC1633-1 as these three highly related plasmids were found in three different species of *Acinetobacter*. Perhaps the rate of conjugative transfer for these plasmids was exceptionally low and below detectable limits in stark contrast to what was reported for pA297-3 in which the conjugative transfer region 1 was uninterrupted. Alternatively, successful conjugative transfer of these plasmids may require certain environmental or media conditions that were not met when the experiments were conducted in the laboratory using established protocols. Further work is clearly needed to resolve this transmissibility conundrum for pAC1530 and pAC1633-1.

**Fig. 6.**
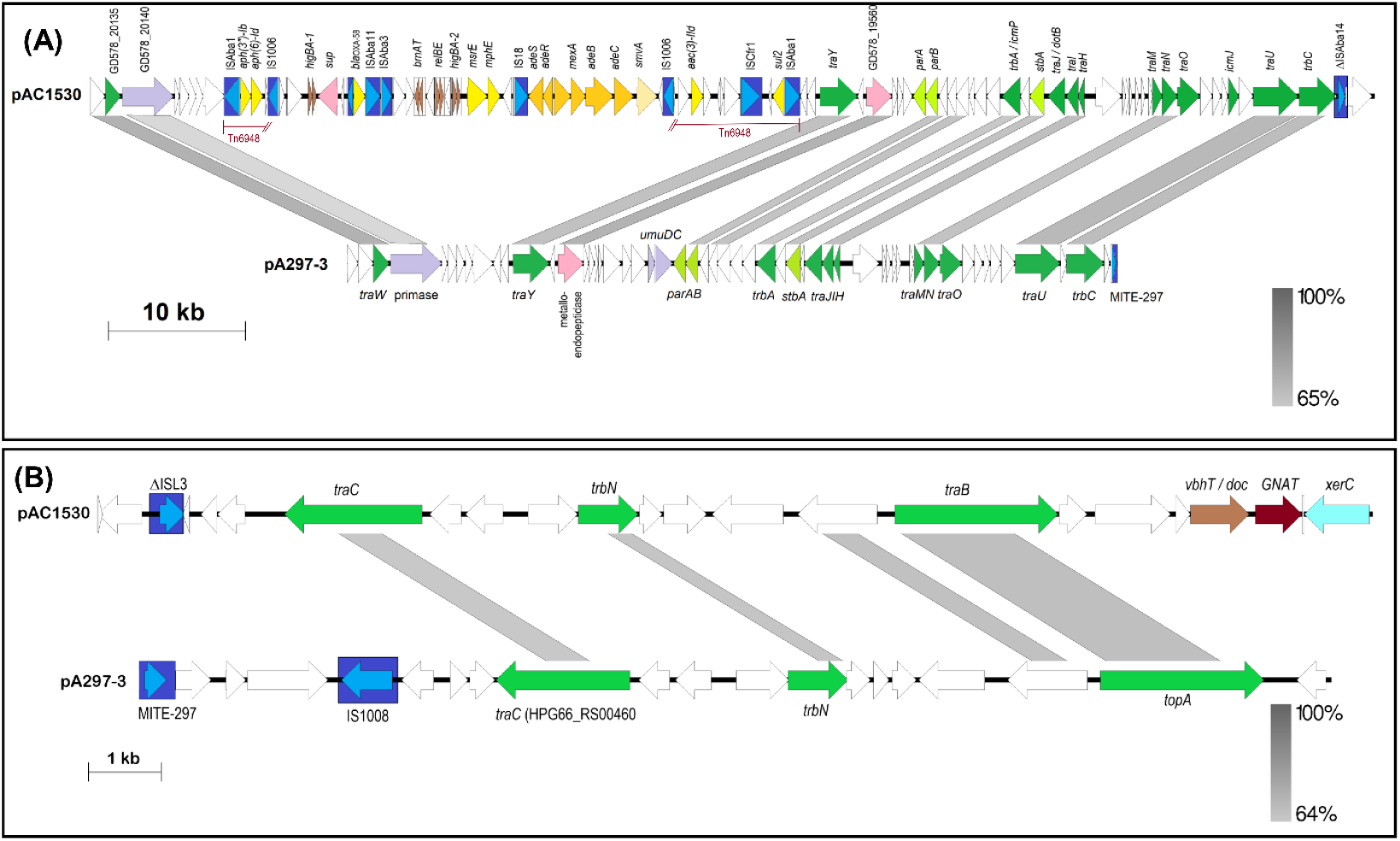
Linearized map of the two conjugative regions of pAC1530 as compared to pA297-3. **(A)** Region 1; map shows nts. 137,720 – 173,972 and continues with 1 – 57,600 of pAC1530 (numbered as in accession no. CP045561.1) and reverse complement of nts. 169,947 – 200,633, continues with 1 – 25,440 of pA297-3 (numbered as in accession no. KU744946.1); **(B)** Region 2; map depicts nts. 80,481 – 98,080 of pAC1530 and reverse complement of nts. 76,001 – 92,455 of pA297-3. Arrows indicate the extents and directions of genes and ORFs with identified conjugative transfer genes depicted as dark green arrows. Lime green arrows are the plasmid partitioning genes *parA* and *parB*. IS elements and the miniature inverted-repeat transposable element (MITE) identified in pA297-3 (72) are depicted as dark blue boxes with their encoded transposase shown as lighter blue arrows. Antibiotic resistance genes in pAC1530 are indicated as yellow arrows while toxin-antitoxin genes are shown as brown arrows. Other identified genes are in purple and pink arrows with white arrows indicating ORFs encoding hypothetical proteins. The IS*Aba1*-flanked Tn*6948* is indicated; note that in pAC1530 and pAC1633-1, Tn*6948* is interrupted by the IS*1006*-flanked region that contains antimicrobial resistance genes such as *bla*_OXA-58_, *sul2*, *msrE* and *mphE* (see **Fig. 5** and main text). Grey-shaded areas indicate regions with DNA sequence identities as indicated by the bars at the bottom right of each figure. Note that although the figure depicts only pAC1530, pAC1633-1 is nearly identical to pAC1530 in Region 1 except for an insertion of IS*Aba11* in the ORF upstream of *traM* and deletion of the *relBE* p*dif* module (see **Fig. 2** and main text).

**Table 3.**
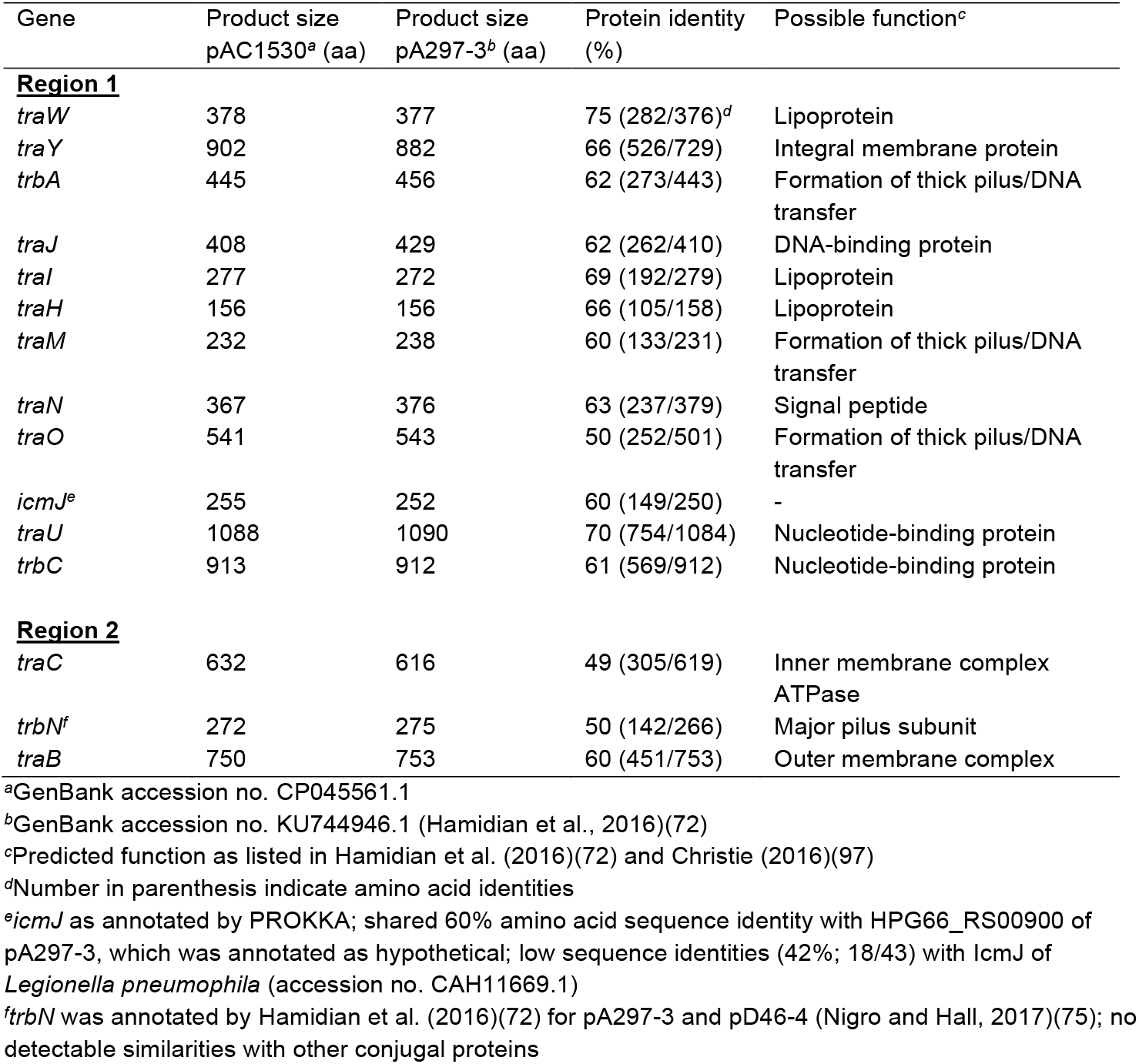
Conjugative transfer-related genes identified in pAC1530 and pAC1633-1 compared to their corresponding genes in pA297-3

### The *tetA(39)* tetracycline resistance gene in pAC1633-2 is within a p*dif* module

Plasmid pAC1633-2 is 12,651 bp and encodes a Rep3 family replicase of the *Acinetobacter* GR8/GR23 group that was preceded by four 22-bp iterons characteristic of Rep3 family plasmids (43, 44). pAC1633-2 also encodes a *tetA(39)* tetracycline resistance gene which was adjacent to and divergently transcribed from a *tetR(39)* regulatory gene **(Fig. 7)**. This 2,001 bp fragment is identical with the *tetAR(39)* genes that made up a p*dif* module in plasmids pS30-1, pRCH52-1 and pAB1-H8 (55). However, the p*dif* sites that flank this *tetAR(39)* region in pAC1633-2 differed from those in pS30-1, pRCH52-1 and pAB1-H8 at the 6-bp spacer and the XerC-recognition site **(Fig. 7)**. Another p*dif* module that was detected in pAC1633-2 encode for the *vapBC* toxin-antitoxin system. Interestingly, when comparing with plasmid pA1296_2 from *A. baumannii* A1296 (accession no. CP018334), the *vapBC* genes flanked by the p*dif* sites were in an inverted orientation (**Fig. 7**), similar to the situation of the *brnTA* toxin-antitoxin p*dif* module in pAC1633-1 and pAC1530 when compared to their similar plasmids. However, the *vapBC* p*dif* modules in these two plasmids were 91% identical in sequence and the sizes of these modules were slightly different: in pAC1633-2, the *vapBC* p*dif* module was 1,259 bp whereas in pA1296_2, it was 1,176 bp.

**Fig. 7.**
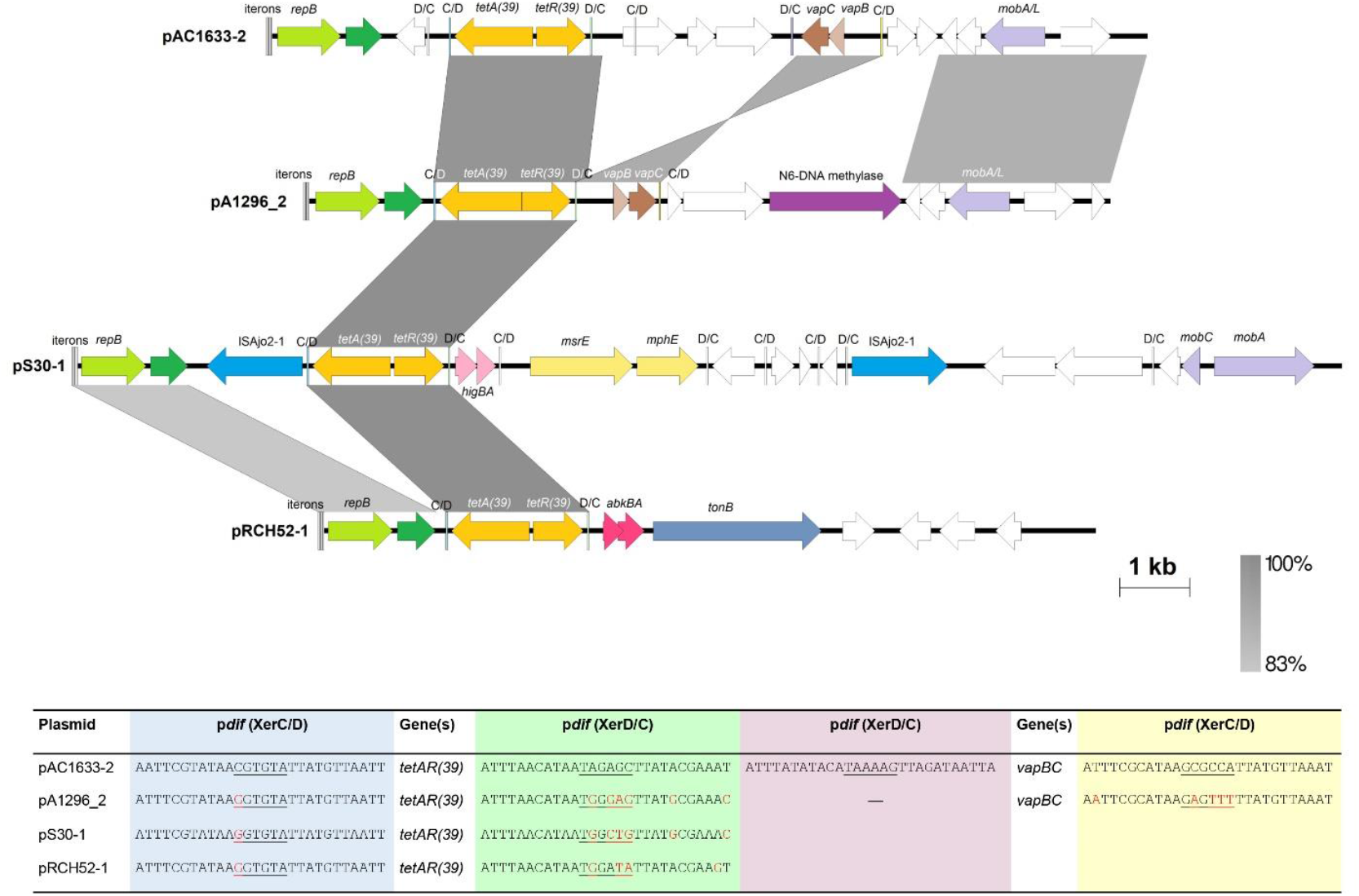
Linearized map of pAC1633-2 compared to other similar plasmids harboring the *tetAR(39)* p*dif* module. Arrows indicate the extents and directions of genes and ORFs with yellow arrows for the *tetA(39)* and *tetR(39)* tetracycline resistance genes and pale yellow arrows for the macrolide resistance genes *msrE* and *mphE* in pS30-1. Plasmid replicase genes of the Rep3 family are depicted as green arrows and labeled *repB* while darker green arrows are ORFs encoding putative DNA-binding proteins that have been previously misannotated as *repA* (44). Mobilization-related genes are shown as light purple arrows and the gene encoding an N6-DNA methylase is in dark purple. The *vapBC* toxin-antitoxin genes in pAC1633-2 and pA1296_2 are indicated as brown arrows; *higBA* in pS30-1 are depicted in pink; whereas *abkBA* in pRCH52-1 are in dark pink. The transposase encoded by ISAjo2 in pS30-1 is colored blue. White arrows are ORFs that encode for hypothetical proteins. Regions with significant DNA identities from 83% - 100% are shaded in shades of grey as indicated in the bar at the bottom right of the figure. The four 22-bp iterons that are the likely *oriV* site for each plasmid is shown as successive horizontal bars at the beginning of the plasmid’s linear map. The p*dif* sites are depicted as horizontal bars labeled as C/D for the XerC-XerD orientation and D/C for XerD-XerC orientation. Note that the orientation of the *vapBC* genes flanked by p*dif* sites in pAC1633-2 are inverted when compared to pA1296_2. The XerC/D and XerD/C sites flanking the *tetAR(39)* genes are colored light blue and light green, respectively, whereas the XerC/D and XerD/C sites flanking the *vapBC* genes in pAC1633-2 are colored light yellow and light purple, respectively. The sequences for the p*dif* sites flanking the *tetAR(39)* genes as well as the *vapBC* TA genes are shown in the table at the bottom of the figure. Bases highlighted in red are those that differ from the p*dif* sequences of pAC1633-2. The accession nos. of the plasmids used in this analysis are as follows: pA1296_2 (accession no. CP018334), pS30-1 (accession no. KY617771) and pRCH52-1 (accession no. KT346360).

pAC1633-2 also harbors a *mobA/L* gene that encodes a relaxase of the MOB_Q_ family, indicating the possibility of the plasmid being mobilized should a suitable conjugative plasmid is present in the host cell. Since AC1633 also harbored the potentially conjugative pAC1633-1 plasmid, the ability of pAC1633-1 to mobilize pAC1633-2 was tested in conjugation experiments by selecting for transconjugants that exhibit tetracycline resistance in addition to azide resistance. Despite repeated experiments, no such transconjugants were detected inferring that pAC1633-1 was likely unable to mobilize pAC1633-2. This, however, does not rule out the possibility that pAC1633-2 could be mobilized by a different type of conjugative plasmid to pAC1633-1.

### The two other plasmids in *A. baumannii* AC1633, pAC1633-3 and pAC1633-4, are cryptic

Two other smaller plasmids are found in *A. baumannii* AC1633, the 9,950-bp pAC1633-3 and the 5,210-bp pAC1633-4. Both plasmids do not encode any antimicrobial, metal resistance or any genes that could confer a specific phenotype to their host and are thus, cryptic plasmids. Both plasmids contain RepB replicases of the Rep3 superfamily that are common in *Acinetobacter* plasmids (43, 44). Comparative analysis of the RepB protein sequences indicate that pAC1633-3 belonged to the recently categorized GR28 group of *Acinetobacter* plasmids (43) whereas pAC1633-4 belonged to the GR7 group. Both pAC1633-3 and pAC1633-4 could potentially be mobilizable as they encode for *mobA/L* genes of the MOB_Q_ family and pAC1633-4 also encode for a *mobS*-like gene (**Suppl. Fig. 6**). However, in the absence of any selectable marker, we were unable to determine if these two plasmids could be mobilized by pAC1633-1.

Four p*dif* sites were detected in pAC1633-3 but none in pAC1633-4. Interestingly, one of the p*dif* modules in pAC1633-3 is a 464 bp region that encodes a putative protein of the SMI1/KNR4 family and is identical with the p*dif* module that was found downstream of the *relBE* p*dif* module in the *A. nosocomialis* AC1530-encoded pAC1530. This SMI1/KNR4 p*dif* module, along with the *relBE* p*dif module*, is absent in pAC1633-1 and is one of the features that differentiated pAC1530 from pAC1633-1. The other p*dif* module in pAC1633-3 is 4,331 bp and encodes a putative regulatory protein of the Xre family, a *hipA*-like toxin, and a 602 amino acid-residues protein of the DEAD/DEAH box family of helicases (**Suppl. Fig. 6**).

## CONCLUSIONS

Complete genome sequencing of carbapenem-resistant *A. baumannii* AC1633 and *A. nosocomialis* AC1530 led to the discovery of a ca. 170 kb plasmid that encoded the NDM-1 and OXA-58 carbapenemases along with several other resistance determinants and was likely responsible for the MDR status of these two clinical isolates. The *A. baumannii* AC1633-encoded pAC1633-1 and the *A. nosocomialis* AC1530-encoded pAC1530 were nearly identical except for the insertion and deletion of IS elements and a p*dif* module. Both plasmids were a patchwork of multiple mobile genetic elements with the *bla*_NDM-1_ residing in a Tn*125* composite transposon while *bla*_OXA-58_ was flanked by IS elements nested within a p*dif* module. The *msrE-mphE* macrolide resistance genes were also located within a p*dif* module, as were several toxin-antitoxin genes, highlighting the importance of these Xer recombination-dependent modules as one of the drivers of plasmid diversity in *Acinetobacter*. Comparative sequence analysis indicated that pAC1633-1/pAC1530 is likely a cointegrate of two plasmids which separately encode the *bla*_NDM-1_ and *bla*_OXA-58_ genes in an *A. pittii* clinical isolate, and that was formed via an IS*1006*-mediated recombination or transposition event. Horizontal transmission of pAC1633-1/pAC1530 was inferred from the discovery of the almost identical plasmid in two different species of *Acinetobacter* from the same hospital but this could not be experimentally demonstrated in the laboratory. Nevertheless, the presence of such large, potentially transmissible multidrug resistant plasmids in *Acinetobacter* that co-harbor the NDM-1 and OXA-58 carbapenemases in this and other recent reports (53, 59) warrants monitoring and assessment of the risk of spread of these plasmids to susceptible strains, particularly in healthcare settings.

## MATERIALS AND METHODS

### Ethical approval, bacterial isolates and antimicrobial susceptibility profiles

Ethical approval for this study was obtained from the Malaysian Ministry of Health’s National Medical Research Register (approval no. NMRR-14-1650-23625-IIR).

*A. baumannii* AC1633 and *A. nosocomialis* AC1530 were isolated from Hospital Sultanah Nur Zahirah, Kuala Terengganu, Malaysia in 2016 and 2015, respectively. Species identification of both isolates was performed by sequencing of the *rpoB* gene as previously described (13, 76). Antimicrobial susceptibility profiles of both isolates were determined using a panel of 22 antibiotics recommended for *Acinetobacter* spp. (26) and by disc diffusion (Oxoid Ltd., Basingstoke, UK) on Mueller-Hinton (MH) agar except for colistin and polymyxin B, which were determined by obtaining the MIC values by the agar diffusion method (25). Carbapenem resistance was validated by determining the MIC values for imipenem, meropenem and doripenem using M.I.C. Evaluator strips (Oxoid Ltd., Basingstoke, UK). Results were interpreted according to the Clinical and Laboratory Standards Institute (CLSI) guidelines (77). Production of metalo-β-lactamases was determined using the Etest MBL kit (bioMérieux, La Balme-les-Grottes, France).

### DNA isolation, whole genome sequencing and sequence analyses

Genomic DNA for whole genome sequencing was prepared using the Geneaid Presto Mini gDNA Bacteria Kit (Geneaid, Taipeh, Taiwan) following the manufacturer’s recommended protocol and the extracted DNA quality was evaluated using a Qubit 2.0 Fluorometer (Life Technologies, Carlsbad, CA). Genome sequencing was performed on the Illumina NextSeq (Illumina Inc., San Diego, CA) and PacBio RSII (PacBio, Menlo Park, CA) platforms by a commercial service provider (Novogene, Beijing, China) and hybrid assembly was carried out using SPAdes (version 3.11.1) (78). Gene prediction for the assembled genomes was performed with PROKKA (79) with annotation achieved using the NCBI Prokaryotic Genome Annotation Pipeline (PGAP) (80). Multilocus sequence typing (MLST) in the Institut Pasteur (81) and Oxford (82) schemes was performed via the *A. baumannii* MLST database (https://pubmlst.org/abaumannii/) (34). Pan genome analysis for *A. baumannii* AC1633, *A. nosocomialis* AC1530 and related global *A. baumannii* and *A. nosocomialis* isolates (as listed in **Suppl. Table 1**) was determined using ROARY with the core genomes identified using the criteria of amino acid sequence identities > 95% (83) and presence in 99% of genomes. The derived core genome alignments for *A. baumannii* and *A. nosocomialis* were then used to infer Maximum-Likelihood (ML) trees using FastTree (84) with 100 bootstraps under the GTR time-reversible model. The resulting *A. baumannii* and *A. nosocomialis* phylogenetic trees were then visualized using iTOL v5 (https://itol.embl.de/) (85).

Antibiotic resistance genes were identified using ResFinder (https://cge.cbs.dtu.dk/services/ResFinder/) (86) and the Comprehensive Antibiotic Resistance Database (CARD) (https://card.mcmaster.ca/) (87) whereas ISFinder (https://isfinder.biotoul.fr/) (42) was used to identify insertion sequences. Toxin-antitoxin systems were identified using the toxin-antitoxin database, TADB 2.0 (https://bioinfo-mml.sjtu.edu.cn/TADB2/index.php) (88), putative conjugative transfer genes were identified using SeCreT4 (https://db-mml.sjtu.edu.cn/SecReT4/) (89). All plasmid sequences were manually inspected using BLAST (https://blast.ncbi.nlm.nih.gov/Blast.cgi) and ORF Finder (https://www.ncbi.nlm.nih.gov/orffinder/) to validate the genes/open reading frames (ORFs) that were predicted by the annotation and other programs. Pfam searches (90) and the NCBI Conserved Domain Database (CDD) (91) were also used to identify possible protein functions. The presence of p*dif* sites in pAC1530, pAC1633-1 and related plasmids was determined by BLASTN screening using known XerC/XerD and XerD/XerC sites in published reports (54–56) and manually examining hits that were 75 – 80% identical in sequence (56).

SnapGene 5.1.5 (GSL Biotech LLC., San Diego, CA) was used to visualize and manipulate the sequences studied. Figures were drawn to scale using EasyFig 2.2.3 (http://mjsull.github.io/Easyfig/) (92) and CGView (http://stothard.afns.ualberta.ca/cgview_server/) (93).

### Conjugation assays

Conjugation assays were carried out to investigate the transmissibility of pAC1530 and pAC1633-1 from their respective carbapenem-resistant natural hosts, *A. nosocomialis* AC1530 and *A. baumannii* AC1633, to the appropriate susceptible isolates, *A. baumannii* ATCC19606 and *A. baumannii* AC1529. *A. baumannii* ATCC19606 is the type strain of *A. baumannii* that has been widely used in various studies, is resistant to sulfonamides due to the presence of the *sul2* gene in its chromosome, but remains susceptible to a wide range of other antibiotics (47, 94) including the carbapenems. *A. baumannii* AC1529 was isolated from the blood of a 59 year old male patient in the Emergency Ward of Hospital Sultanah Nur Zahirah in 2015 and its identity was confirmed by *rpoB* sequencing (76). AC1529 showed intermediate resistance to cefotaxime and ceftriaxone but was susceptible to the other 20 antimicrobials that were tested including carbapenems. *A. baumannii* ATCC19606 and *A. baumannii* AC1529 were selected for induction to azide resistance to be used as recipient strains in the conjugation assays. Spontaneous mutation of both *A. baumannii* strains to sodium azide resistance was performed by continuous exposure to increasing concentrations of sodium azide as described by Leungtongkam et al. (95).

*A. baumannii* ATCC19606 and *A. baumannii* AC1529 isolates with sodium azide MIC values > 300 μg/ml were used as recipients whereas *A. baumannii* AC1633 and *A. nosocomialis* AC1530 were used as donors in four separate conjugation experiments. Equal amounts of overnight cultures of the donor and recipient cells were mixed and incubated at 37°C on Luria-Bertani (LB) agar plates overnight. Cells were resuspended and diluted in 0.9% NaCl and selected on LB agar plates supplemented with 300 μg/ml sodium azide and 10 μg/ml imipenem. Conjugation assays were also repeated with different ratios of donor to recipient cells (1:2, 1:3, 2:1 and 3:1). To investigate if the *tetA(39)*-harboring plasmid pAC1633-2 could be mobilized by pAC1633-1 in *A. baumannii* AC1633, conjugation experiments involving AC1633 as donor were also plated on LB agar plates supplemented with 300 μg/ml sodium azide and 4 μg/ml tetracycline.

### Accession nos

The complete sequence of the *A. baumannii* AC1633 chromosome was deposited in GenBank under accession no. CP059300 whereas its four plasmids were deposited under the following accession nos.: pAC1633-1 (CP059301), pAC1633-2 (CP059303), pAC1633-3 (CP059304) and pAC1633-4 (CP059302). The *A. nosocomialis* AC1530 chromosomal sequence was deposited under accession no. CP045560.1 whereas its plasmid pAC1530 was deposited under accession no. CP045561.1.

## ACKNOWLEDGEMENTS

We thank Dr Fatimah Haslinda Abdullah and Dr Norlela Othman of Hospital Sultanah Nur Zahirah, Kuala Terengganu, for supplying the AC1530 and AC1633 isolates used in this study. This study was funded by provisions of the following grants: Fundamental Research Grant Scheme **FRGS/1/2017/SKK11/UNISZA/02/4** from the Malaysian Ministry of Higher Education and University Laboratory Materials Grant **UniSZA/LABMAT/2018/09** from Universiti Sultan Zainal Abidin.

