## Supplementary Data for "Complete genome sequencing of *Acinetobacter baumannii* AC1633 and *Acinetobacter nosocomialis* AC1530 unveils a large multidrug resistant plasmid encoding the NDM-1 and OXA-58 carbapenemases"

### SUPPLEMENTARY FIGURES AND TABLES

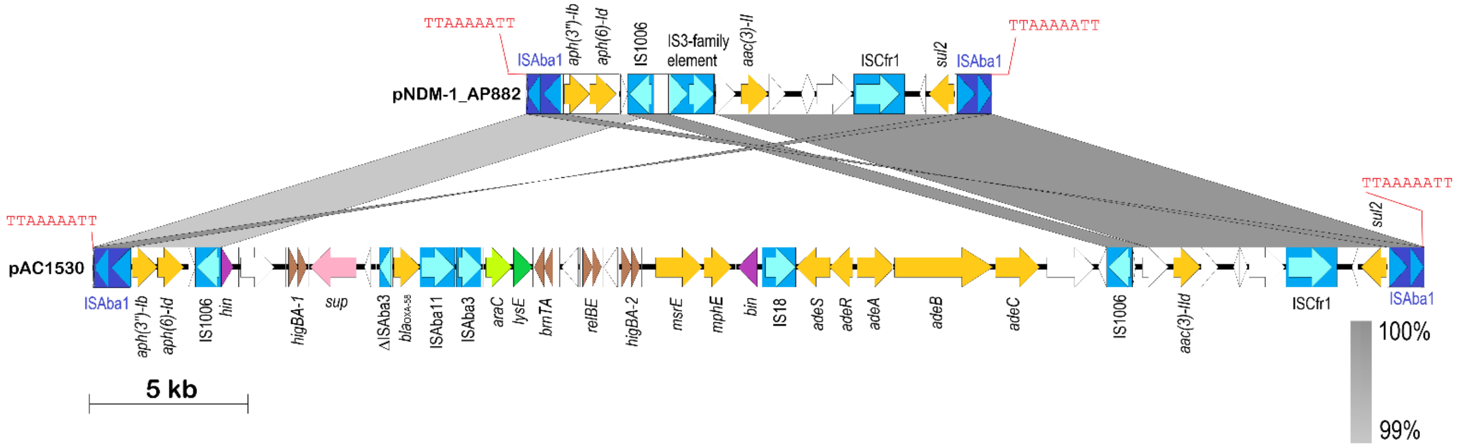


**Suppl. Fig. 1: Linear map of the IS*Aba1*-flanked composite transposon Tn*6948* in the *A. pittii* AP882-encoded pNDM-1_AP882 in comparison with the transposon in pAC1530.** Arrows indicate extents and directions of genes and ORFs. Insertion sequence (IS) elements are depicted as blue-colored boxes with lighter blue arrows showing the directions of the transposase genes. IS*Aba1* that flanks the composite transposon is depicted in darker shades of blue. The 9-bp direct repeat sequences that flank the composite transposon is shown in red fonts. Antimicrobial resistance genes are shown as yellow arrows; brown arrows are toxin-antitoxin systems; purple arrows are putative DNA recombination genes; other colored arrows are genes with known homologs; white arrows are ORFs encoding hypothetical proteins. Grey-shaded areas indicate regions with >99% nucleotide sequence identities. The linear map of pNDM-1_AP882 covered nts. 146,571 – 146,597 and continued with nts. 1 – 14,741 of accession no. CP014478 whereas the map of pAC1530 covered nts. 147,515 – 173,972 and continued with nts. 1 – 15,685 of accession no. CP045561.1. Note that plasmid pAC1633-1 (accession no. CP059301) is almost identical to pAC1530 within this region except for the omission of the *relBE* toxin-antitoxin system and its downstream ORF as detailed in the text and in Fig. 4.

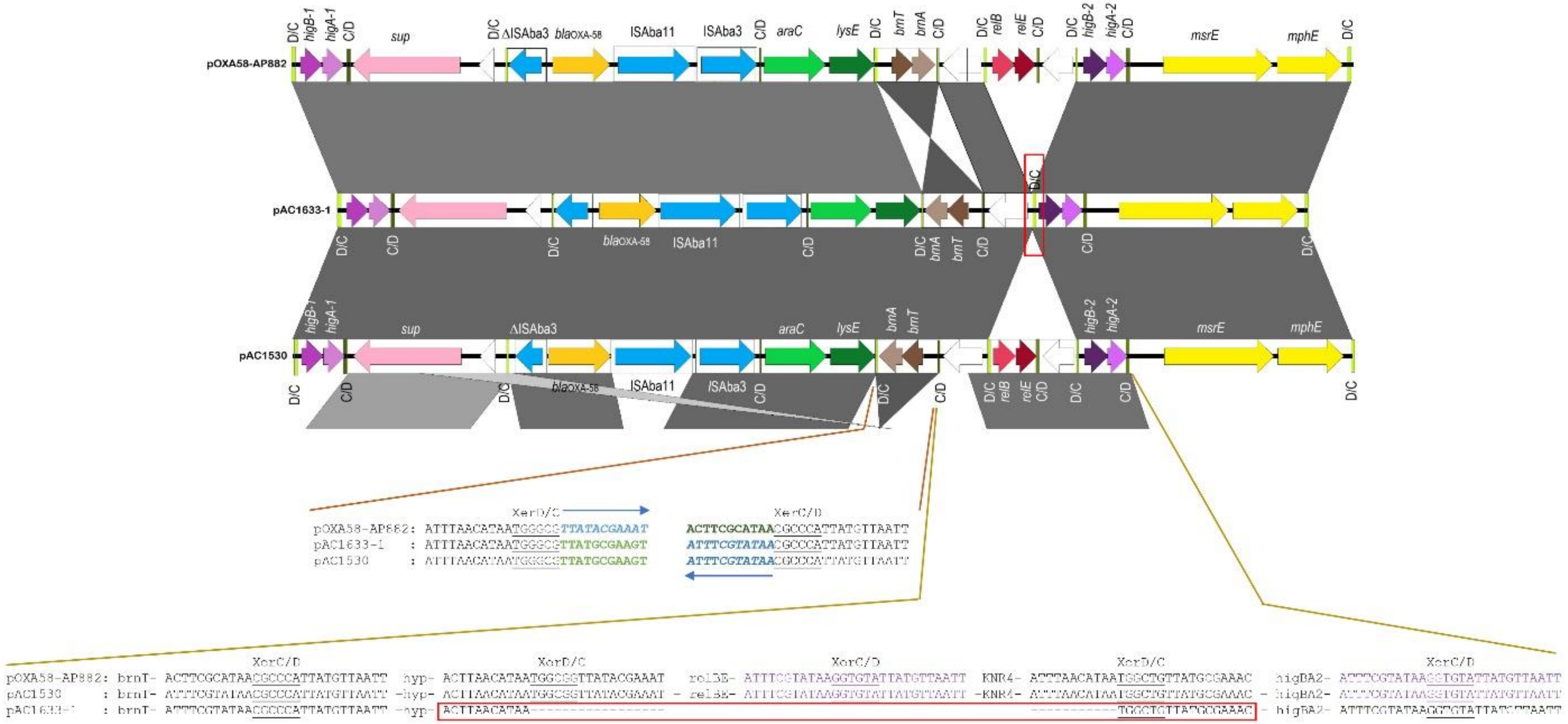


**Suppl. Fig. 2: Comparison of p*dif* sequences in pAC1633-1/pAC1530 and pOXA-58_AP882 at the inversion region encompassing *brnTA* and the insertion point of the *relBE* TA system.** The comparative map and coordinates used for pOXA-58_AP882, pAC1633-1 and pAC1530 are as in Fig. 4. The p*dif* sequences (either XerD/C or XerC/D) are indicated for the respective plasmids and the relevant regions. For the *brnTA* p*dif* module, note that the XerC sequences in pOXA-58_AP882 are in reverse orientation when compared to the corresponding XerC sequences in pAC1633-1 and pAC1530 and these are indicated by blue arrows as well as blue and green fonts for the respective p*dif* XerC site. Also note that the p*dif* sequences downstream of the *relE* gene are identical with the p*dif* sequences downstream of the *higA-2* gene, and these sequences are in purple fonts. The p*dif* (XerD/C) site in pAC1633-1 in which the *relBE*-SMI1/KNR4 p*dif* module was inserted in pAC1530 is outlined in a red box and its sequences outlined likewise below. Note that for this particular p*dif* site, the XerD sequences are identical to the XerD sequences of the p*dif* site upstream of *relB* while the 6-bp spacer and XerC sequences are identical to the p*dif* site upstream of *higB-2*.

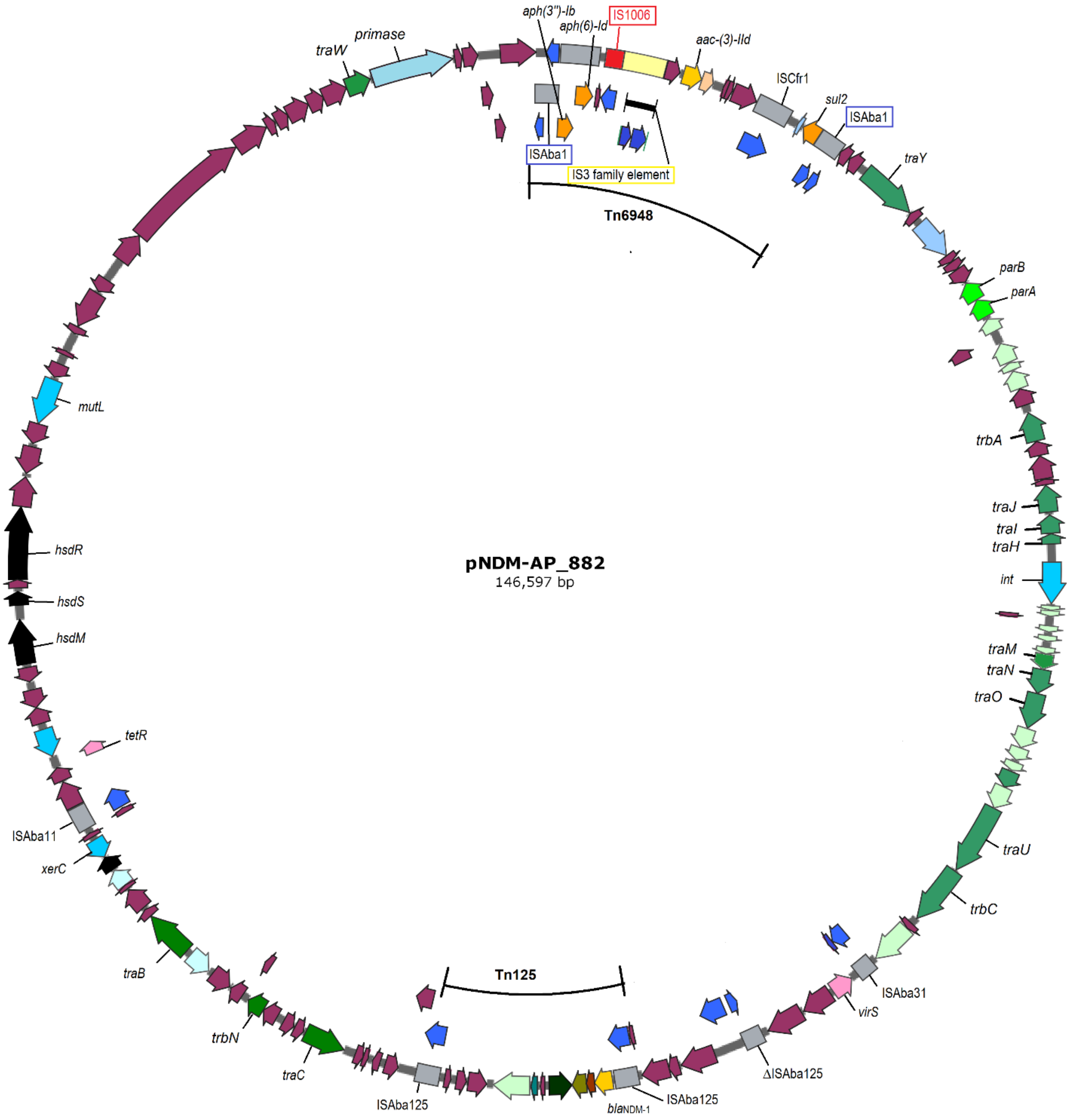


**Suppl. Fig. 3. Circular map of plasmid pNDM-1_AP882 from *Acinetobacter pittii* AP882 (accession no. CP014478).** The single IS*1006* copy is indicated in red. The 1,940 bp region adjacent to IS*1006* that differed to pAC1530/pAC1633-1 and contained an IS-like element of the IS*3* family is indicated in a pale-yellow box. The two IS*Aba1* elements that flanked the composite transposon Tn*6948* is highlighted in blue boxes. Antimicrobial resistance genes including *bla*$_{\mathrm{NDM-1}}$ is shown as gold and orange arrows, other IS elements are as grey boxes with their encoded transposases in blue arrows. Green arrows are genes/ORFs that are likely involved in conjugative gene transfer with darker green arrows showing genes that have homology with known conjugative transfer genes. Sky blue arrows are genes that are involved in DNA recombination and repair. Maroon arrows are ORFs/CDS encoding for hypothetical proteins.

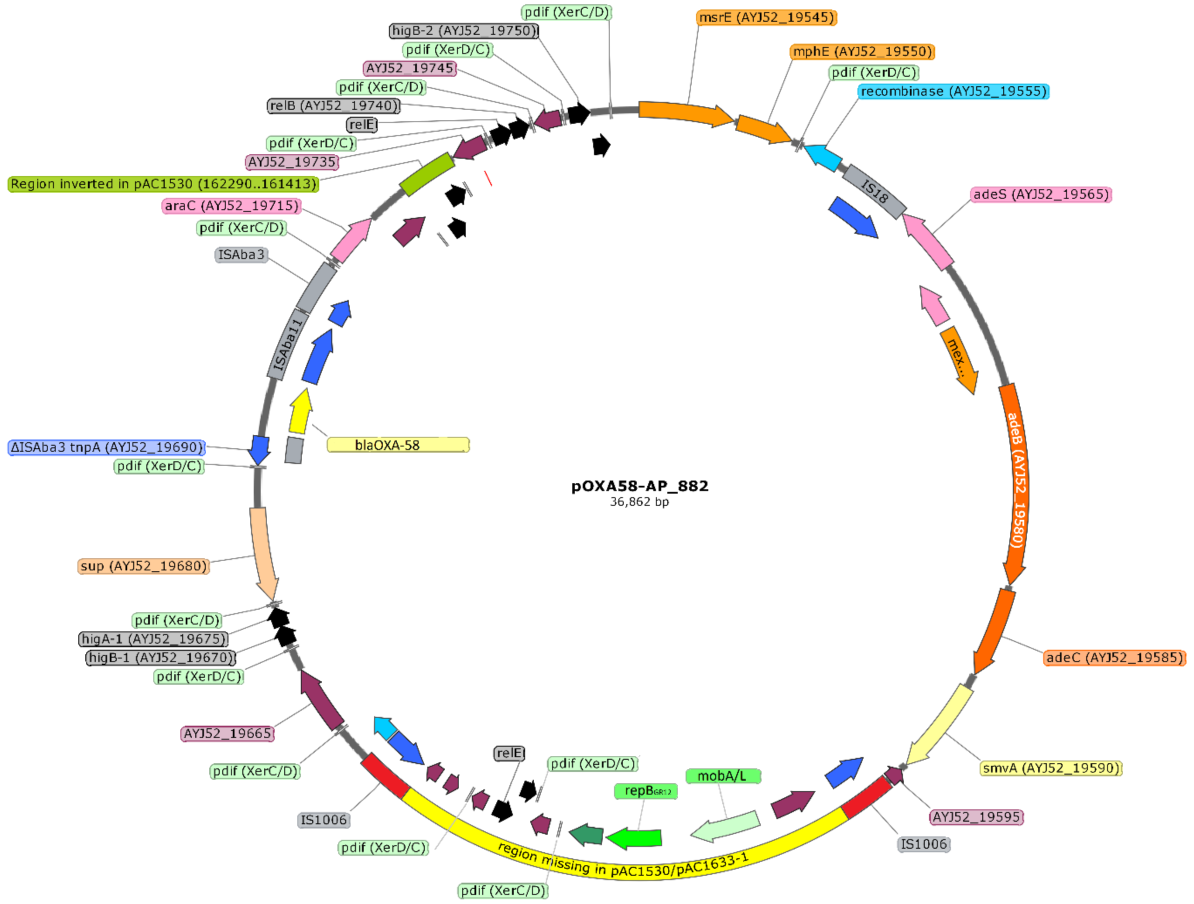


**Suppl. Fig. 4. Circular map of plasmid pOXA-58_AP882 from *Acinetobacter pittii* AP882 (accession no. CP014479) showing distribution of p*dif* sites.** The two copies of IS*1006* are depicted in red, IS-encoded transposases are shown in blue arrows. The 7,191 bp fragment of pOXA-58_AP882 that is absent from pAC1530 and pAC1633-1 is labeled and highlighted in yellow. This region contained the *mobA/L* gene, *repB*$_{\text{GR12}}$ encoding a Rep3 family plasmid replicase of the *Acinetobacter* GR12 family, an ORF encoding a helix-turn-helix putative DNA-binding protein depicted as a dark green arrow downstream of *repB*$_{\text{GR12}}$, and a putative *relE-xre* toxin-antitoxin system shown as black arrows. Other TA systems are also indicated as black arrows and labeled accordingly. p*dif* sites are depicted as horizontal bars and are labeled as per the orientation of their XerD- and XerC-binding sites (i.e., either XerC/D or XerD/C). The *bla*$_{\text{OXA-58}}$ gene is shown as a yellow arrow and labeled.

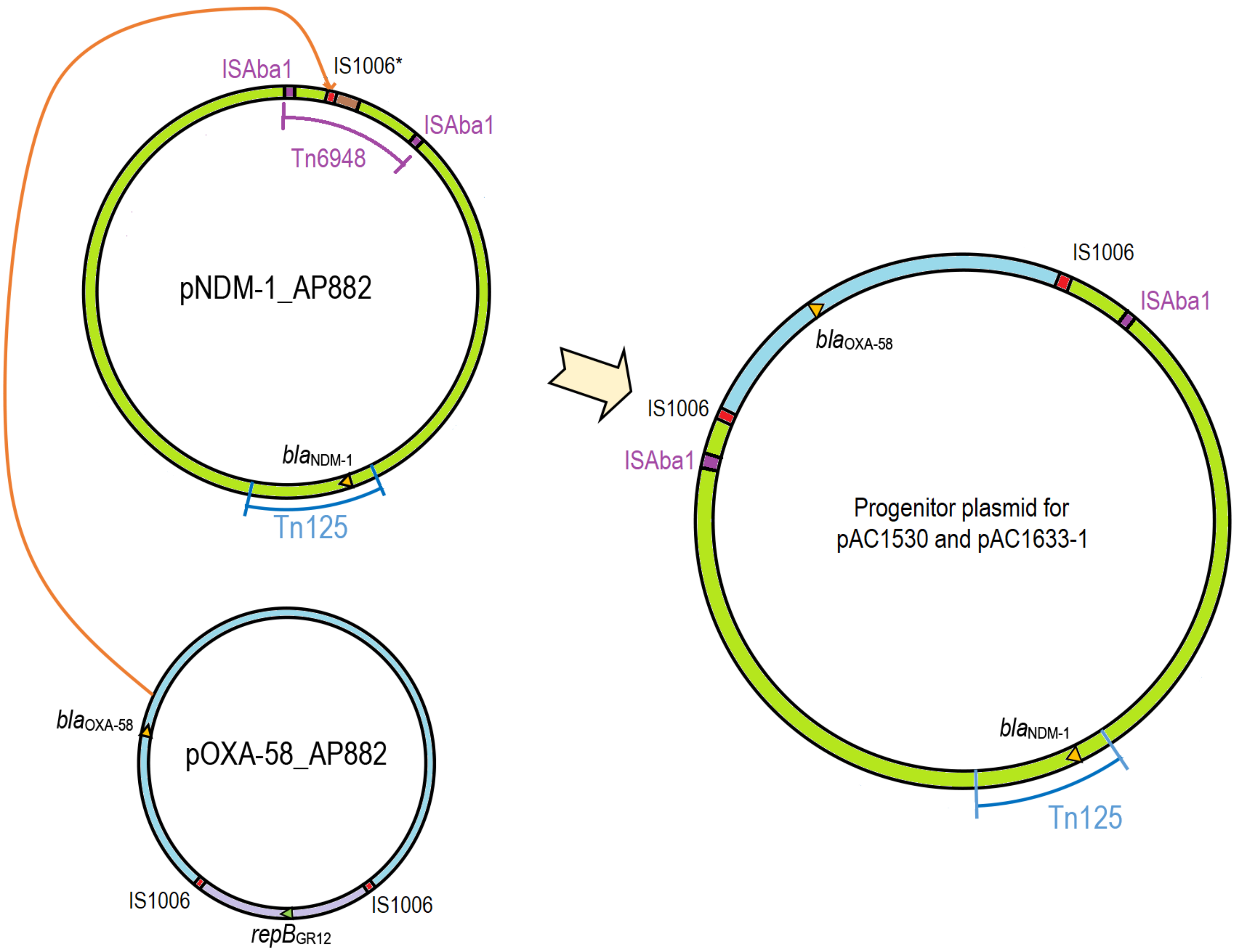


**Suppl. Fig. 5. Hypothetical IS*1006*-mediated cointegration for the generation of the predecessor plasmid for pAC1530 and pAC1633-1 from pNDM-1_AP882 and pOXA-58_AP882.** The 29,671 bp region of pOXA-58_AP882 which contained the *bla*$_{OXA-58}$ gene and is flanked by IS*1006* (labeled in light blue) likely recombined with pNDM-1_AP882 at its single IS*1006* region leading to the formation of the cointegrate predecessor plasmid for pAC1530 and pAC1633-1 that harbored both *bla*$_{OXA-58}$ and *bla*$_{NDM-1}$. The light purple region in pOXA-58_AP882 contains the *repB* replicase gene of the *Acinetobacter* GR12 family and was not involved in the plasmid cointegrate formation. IS*1006* is depicted as red boxes and labeled. pNDM-1_AP882 harbored an IS*1006* with a T175C mutation (when compared with the IS*1006* sequences in pOXA-

58_AP882) and is labeled as IS*1006**. The brown-colored box in pNDM-1_AP882 refers to a 1,940 bp region adjacent to IS*1006** that was absent in pAC1530 and pAC1633-1 and contained an IS*3* family element. This region could have been deleted during the IS*1006*-mediated cointegration process or after the formation of the cointegrate plasmid. The IS*Aba1* elements that flanked the composite transposon Tn*6948* in pNDM-1_AP882, is indicated in purple boxes and labeled.

(A)

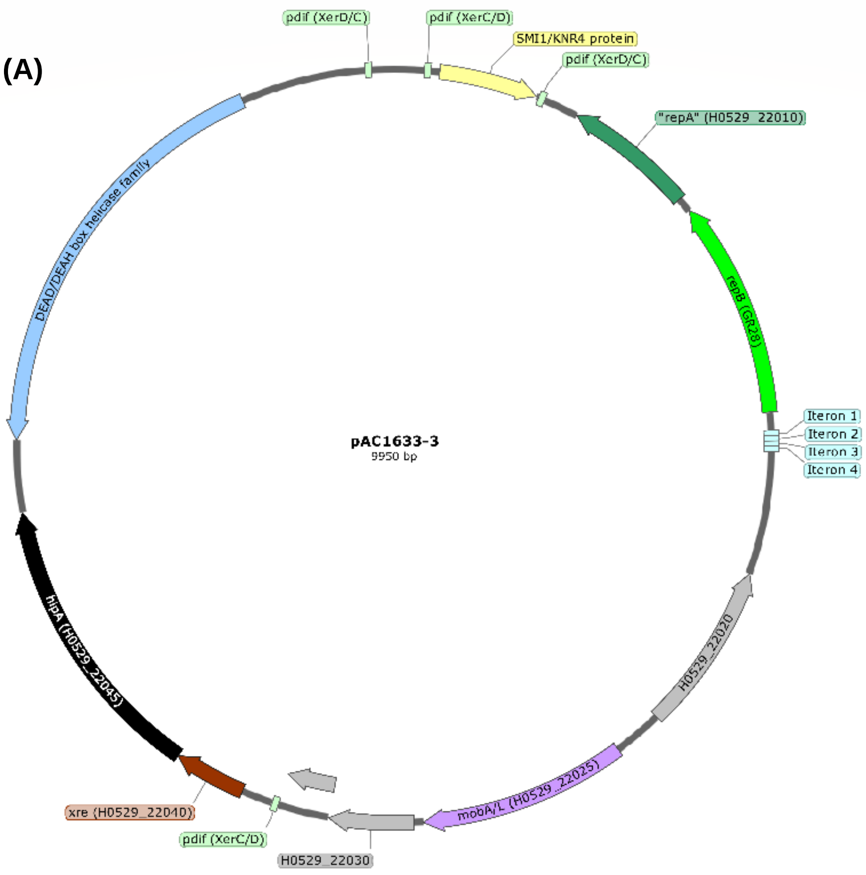


(B)

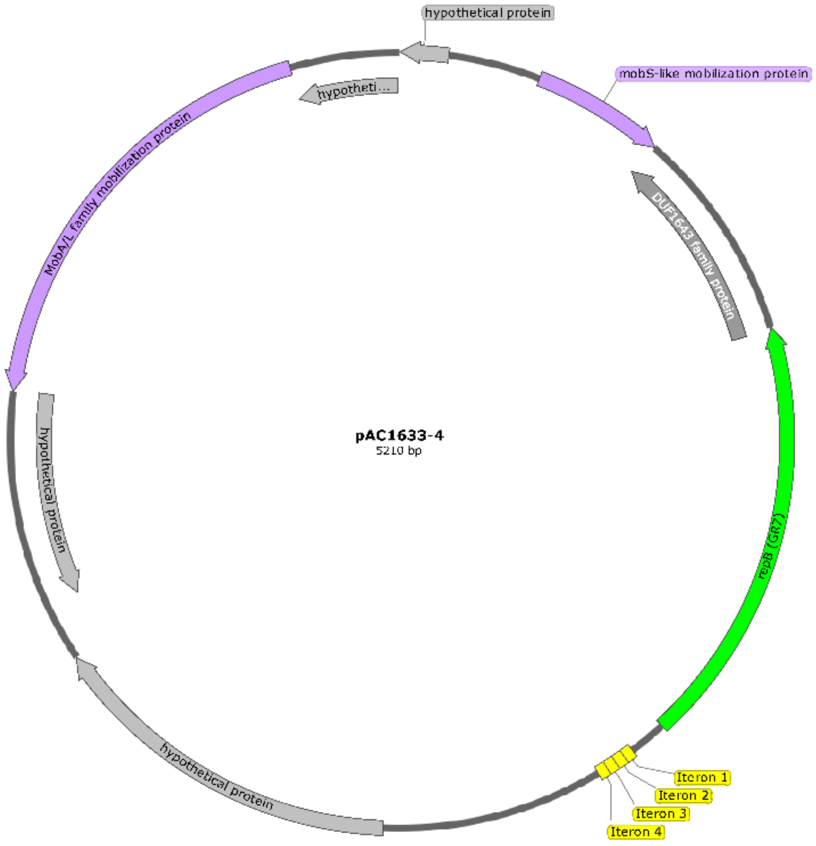


**Suppl. Fig. 6. Map of the small cryptic plasmids in *A. baumannii* AC1633. (A)** Circular map of pAC1633-3; **(B)** circular map of pAC1633-4. The *repB* gene encoding the plasmid replicase in both plasmids is indicated as a green arrow with their *Acinetobacter* plasmid GR grouping number labeled in parentheses; the darker green arrow in pAC1633-3 **(A)** refers to the putative DNA-binding protein encoded by the ORF downstream of *repB* that has been commonly misannotated as *repA* (1). The four 22-bp iterons that are likely the origin of replication for the plasmids are shown as blue rectangles in pAC1633-3 **(A)** and yellow rectangles in pAC1633-4 **(B)** to indicate the difference in iteron sequences. Genes that are associated with plasmid mobilization are depicted as light purple arrows. The *hipA* toxin gene in pAC1633-3 is indicated in black arrow whereas its putative antitoxin/regulator gene, *xre*, is shown in brown arrow upstream of *hipA*. The p*dif* sites are indicated as light green rectangles and labeled accordingly. The gene encoding a putative SMI1/KNR4 family protein that encompassed a 464 bp p*dif* module that was also present in pAC1530 but absent in pAC1633-1 is depicted as a light-yellow-colored arrow. No p*dif* sites could be detected in pAC1633-4. Grey arrows refer to ORFs encoding hypothetical proteins.

**Suppl. Table 1A. Characteristics of the *Acinetobacter baumannii* strains that were used in comparative genomics and phylogenetic tree construction in Fig. 1A**

| Isolate | Source of isolate/ Location/ Year[a] | MLST | | Clonal Group | Accession nos. / Reference(s) |
|---|---|---|---|---|---|
| | | Oxford | Pasteur | | |
| A85 | Sputum / Australia / 2003 | 781[Δ] | 1 | GC1 | CP021782; (2, 3) |
| A388 | ND / Greece / 2002 | 439 | 1 | GC1 | CP024418.1; (4) |
| AB0057 | Blood /Walter Reed Army Hospital, Washington / 2004 | 207 | 1 | GC1 | CP001182; (5) |
| AB307-0294 | Blood / New York / 1994 | 231 | 1 | GC1 | CP001172.2; (3) |
| AC12 | Blood / Kuala Terengganu / 2011 | 195 | 2 | GC2 | CP007549; (6) |
| AC29 | Blood / Kuala Terengganu / 2011 | 195 | 2 | GC2 | CP007535; (7) |
| AC30 | Blood / Kuala Terengganu / 2011 | 195 | 2 | GC2 | CP007577; (7) |
| ACICU | CSF / Rome / 2005 | 437 | 2 | GC2 | CP031380; (3, 8) |
| ATCC17978 | Blood / ND / 1951 | 112 | 437 | | CP012004.1; (9, 10) |
| ATCC19606 | Urine / USA / 1948 (type strain) | 931 | 52 | | CP046654.1; (11) |
| AYE | ND / France / 2001 | 231, 1604[#] | 1 | GC1 | CU459141.1; (12, 13) |
| BJAB0715 | CSF / Beijing / 2007 | 642 | 23 | | CP003847.1; (14) |
| BJAB0868 | Ascites / Beijing / 2008 | 218* | 2 | GC2 | CP003849.1; (14) |
| BJAB07104 | Blood / Beijing / 2007 | 368, 1962* | 2 | GC2 | CP003846,1; (14) |
| Canada-BC-5 | ND / Canada / 2006 | 947[§] | 1 | GC1 | AFDN00000000; (15) |
| CIP70.10 | Skin tissue/ France/1970 | 819 | 126 | | LN865143; (16) |
| D1279779 | Blood / Darwin, Australia / 2009 | 942 | 267 | | CP003967.2; (17) |
| IS-123 | Wound / Baghdad, Iraq / 2009 | 928 | 3 | GC3 | ALII00000000; (15) |
| K50 | Urine / Kuwait / 2008 | 499 | 158 | | OHJL00000000.1; (18) |
| LAC-4 | ND /Los Angeles/1997 | 447 | 10 | | CP007712.1; (19) |
| M1 | ND/ Malaysia / 2009 | 195 | 2 | GC2 | LAIL00000000.1 |
| MDR-TJ | ND / Tianjin / ND | 369, 1837* | 2 | GC2 | CP003500.1; (20, 21) |

| | | | | | |
|---|---|---|---|---|---|
| MDR-ZJ06 | Blood/ Hangzhou / 2006 | 643* | 2 | GC2 | CP001937.2; (22) |
| Naval-81 | Blood / Betheseda, USA / 2006 | 928 | 3 | GC3 | AFDB00000000; (15) |
| OIFC098 | ND / Germany / 2003 | 391 | 10 | | AMDF00000000; (15) |
| OIFC137 | Catheter tip / Washington / 2003 | 106 | 3 | GC3 | AFDK0000000; (15) |
| PR07 | Blood / Malaysia / ND | 734 | 239 | | CP012035.1; (23) |
| R2090 | Rectal swab / Egypt / ND | 942 | 267 | | LN868200.1; (24) |
| RBH3 | Endotracheal aspirate / Australia / 2002 | 781$^{\Delta}$ | 1 | GC1 | FBXD00000000; (4) |
| SDF | Human body louse / ND/ ND | NA | 17 | | CU468230.2; (13) |
| TCDC | Blood / Taiwan / ND | 218* | 2 | GC2 | CP002522.2; (25) |
| TYTH-1 | Blood / Taiwan / 2006 | 455* | 2 | GC2 | CP003856.1; (26) |
| WC-A-694 | ND / Washington / 2008 | 928 | 3 | GC3 | AMTA00000000; (15) |
| ZW85-1 | Feces / China / ND | 378 | 639 | | CP006768.1; (27) |
| 341 | Sputum / Malaysia / 2013 | 938 | 2 | GC2 | JQSD00000000 |
| 461 | Wound swab / Malaysia / 2013 | 195 | 2 | GC2 | LCTE00000000 |
| 863 | Sputum / Malaysia / 2014 | 938 | 2 | GC2 | LZTF01000000 |
| 1656-2 | Sputum / South Korea / 2004 | 423 | 2 | GC2 | CP001921.1; (28) |
| 9102 | Bronchial fluid/ Mexico/ ND | 231 | 1 | GC1 | CP023029.1; (29) |
| 5845 | Wound/ Mexico/ 2009 | 417 | 2 | GC2 | NZ_CP023034.1; (29) |
| 10042 | Secretion/ Mexico/ 2011 | 473 | 2 | GC2 | NZ_CP023026.1; (29) |

[a] ND – unknown

* Two *gdh-B* alleles found (*gdh-B-3* and *gdh-B-189*) which could correspond to more than one STs

[#] Two *gdh-B* alleles found (*gdh-B-4* and *gdh-B-162*) which could correspond to more than one STs

[§] Two *gdh-B* alleles found (*gdh-B-74* and *gdh-B-182*)

[Δ]Two *gdh-B* alleles found (*gdh-B-4* and *gdh-B-182*)

#### Suppl. Table 1B. Characteristics of the *Acinetobacter nosocomialis* strains that were used in comparative genomics and phylogenetic tree construction in Fig. 1B

| Isolate | Source of isolate / Location / Year[a] | MLST | | Accession nos. / Reference(s) |
|---|---|---|---|---|
| | | Oxford | Pasteur | |
| 2010S01-197 | Respiratory/Taiwan/2010 | 1999* | 1272* | CP033561.1; (30) |
| 2010N17-248 | Blood/Taiwan/2010 | 1996* | 410 | CP033572.1; (30) |
| 2012C01-137 | Blood/Taiwan/2012 | 1343* | 217* | CP033557.1; (30) |
| 2014N23-120 | Blood/Taiwan/2014 | 1996* | 410* | CP033545.1; (30) |
| 2014S01-097 | Respiratory/Taiwan/2014 | 1996* | 410* | CP033550.1; (30) |
| 28F | ND/ Colombia /ND | 948 | 71 | CBSD000000000.2; (31) |
| 6411 | ND/Colombia/2012 | 1162 | 322 | CP010368 |
| Ab22222 | ND | 1066 | 71 | AKAR01000001 |
| AB6 | Blood/Guanzhou/2013 | 708 | 782 | PXNE01000001 |
| AB7 | Blood/Guanzhou/2014 | 708 | 782 | PXND01000001 |
| AB11 | Blood/Guanzhou/2014 | 2078 | 1264 | PXMZ01000001.1 |
| FDAARGOS-129 | Abscess/USA/2014 | 958 | 433 | CP014019.1; (32) |
| HJ14 | Kidney fluid/Zhejiang/2014 | 1343 | 217 | MADF01000001.1 |
| J1A | Sea water/Phillipine Sea/2016 | 715 | 768 | CP042994.1 |
| KAN01 | Sputum/Daejon, Korea/2015 | 1740 | 768 | CP038816.1 |
| KAN02 | Blood/Jeonju, Korea/2015 | 2078 | 1264 | CP036171.1 |
| LMG10619 | Sputum/Japan/ND | 1035 | 76 | BBSR01000001.1 |
| M2 | Hip infection /Ohio, USA/1996 | ND | ND | CP040105.1; (33, 34) |
| NCTC8102 | ND/Rhode Island, USA/1950 | 958 | 74 | CP029351.1; (35) |
| NIPH386 | Sputum/Czech Republic/1996 | 948 | 410 | KB849561.1 |
| NIPH2119 | Sputum/Rotterdam/1987 | 1035 | 76 | APOP01000001.1 |
| P020 | ND/Taiwan/2017 | 2078 | 1264 | APCE01000001.1 |
| SSA3 | Blood/Seoul/2013 | 958 | 433 | CP020588.1; (36) |
| T228 | ND/Bangkok/2010 | 1897 | 279 | JRUA01000001.1; (37) |
| UBA6823 | Metal/New York/ND (Metagenome sample) | 854[#] | 1259* | DKEL01000001.1; (38) |

[a] ND – unknown
* alleles with less than 100% identity/imperfect hit
[#] Nearest ST

**Suppl. Table S2. Carriage of efflux-mediated antimicrobial resistance genes in *A. baumannii* AC1633 and *A. nosocomialis* AC1530**

| Strain | Efflux pump family | | |
|---|---|---|---|
| | Resistance-nodulation division (RND) family[a] | Major facilitator superfamily (MFS) | Small multidrug resistance (SMR) family |
| *A. baumannii* AC1633 | | | |
| • Chromosome | • *adeI, adeJ, adeK*, *adeN*, *adeF*, *adeG*, *adeH*, *adeL* *adeR* | • *abaF, abaQ, amvA* | • *abeS* |
| • pAC1633-1 | • *adeR, adeS, adeA, adeB, adeC* | | |
| • pAC1633-2 | | • *tetA(39)* | |
| • pAC1633-3 | | | |
| • pAC1633-4 | | | |
| *A. nosocomialis* AC1530 | | | |
| • Chromosome | • *adeF, adeG, adeH, adeL, adeR* | • *amvA, abaQ* | • *abeS* |
| • pAC1530 | • *adeR, adeS, adeA, adeB, adeC* | | |

[a] Genes in the same colored font indicate that they are part of an operon
